# The minimal cell-cycle control system in *Marchantia* as a framework for understanding plant cell proliferation

**DOI:** 10.1101/2025.03.12.642684

**Authors:** Facundo Romani, Ignacy Bonter, Marius Rebmann, Go Takahashi, Fernando Guzman-Chavez, Francesco De Batté, Yuki Hirakawa, Jim Haseloff

## Abstract

The regulation of cell division is broadly conserved across eukaryotes, governed by cyclins and cyclin-dependent kinases (CDKs) to coordinate progression through the cell cycle. Plants have evolved a complex set of cell-cycle genes with unique features. The high number of cyclin-CDK pairs in flowering plants complicates functional studies due to redundancy and diversification. It is critical to study simple systems in other plant lineages to better understand the functional integration of the cell-cycle control machinery and its evolution across land plants.

Through a comprehensive phylogenetic analysis, we show that non-seed plants possess a simple repertoire of cyclin and CDK proteins, suggesting that the observed complexity in seed plants is a derived trait. The liverwort *Marchantia polymorpha* possesses a streamlined set of core cell cycle genes with minimal redundancy during vegetative development. Using single-cell RNA-seq and fluorescent reporters, we found a precise, phase-specific expression pattern for cell cycle genes. We demonstrated *in vivo* that only three cyclins are active, one at a given phase, without redundancy. Functional analyses revealed that Mp*CYCD;1* promotes cell cycle re-entry and disrupts differentiation, while overexpression of Mp*CYCA* or Mp*CYCB;1* arrests the cell cycle, consistent with their respective roles at G1, S, and G2/M progression.

Our findings highlight the functional conservation of mechanisms for cell-cycle control across eukaryotes and provide insights into its ancestral state, revealing a minimal set of functional components required for multicellular development. This study advances our understanding of fundamental aspects of cell-cycle regulation and opens new possibilities for engineering plant growth.

## INTRODUCTION

Eukaryotic organisms have evolved sophisticated mechanisms to control cell proliferation. While cell-cycle control shares common features across eukaryotes, with a conserved set of molecular mechanisms, different lineages such as plants, animals, and fungi adapted it to different multicellular body plans and developmental programmes. Understanding these regulatory networks is a central question for developmental biology and the evolution of development. This knowledge could be instrumental for rational engineering of development and neoorganogenesis in crop plants and promote advances in agricultural biotechnology (Brophy et al., 2018).

The eukaryotic mitotic cell cycle progress through G1 (gap 1), S (synthesis), and G2 (gap 2) phases to in mitosis (M) (Mironov et al., 1999; Krylov et al., 2003). The transition between phases is governed by a relay of cyclins that activate cyclin-dependent kinases (CDKs) to ensure the proper coordination of replication and chromosome segregation. Group I cyclins are unequivocally associated with cell cycle regulation (Martinez-Alonso and Malumbres, 2020), following an archetypical sequence: D-type cyclins regulate the G1/S transition, A-type cyclins are linked to the S-phase, and B-type cyclins with the G2/M transition (Inze and De Veylder, 2006; Komaki and Sugimoto, 2012). The interaction of cyclins with their CDK partners phosphorylate the retinoblastoma (RB) protein and trigger the transduction of signals necessary for the division of the cells. The regulation of these networks is not only crucial for controlling the rate of cell division but also for maintaining the appropriate balance between proliferation and differentiation, and essential for functional integrity, proper organ formation and integration of environmental responses (Inze and De Veylder, 2006).

While the canonical organization of phase specific cyclins/CDKs is well portrayed in the literature and textbooks, most model organisms have different sets of genes that further complicate and specialize this system. For example, animals have E-type cyclins, another G1 specific cyclin that overlaps with D-type, and yeast do not have a true A-type cyclin ortholog. This question is of special importance in plants. The first plant cell cycle components were identified in *Arabidopsis thaliana* as homologs of yeast and animal regulators (Gutierrez, 2009). Plants also exhibit unique cell cycle features, such as a plant-specific CDKB expressed during G2/M (Gutierrez, 2009). The high complexity of cyclin and CDK regulation, especially in flowering plants, has made it challenging to disentangle their precise functional roles in the progression of the cell cycle in plants (Gutierrez, 2022) and has hindered our ability to understand many fundamental aspects of the cell cycle regulation and evolution.

The Arabidopsis genome displays a large repertoire of cell cycle genes, including 31 Group I cyclins. Among the 10 D-type cyclins, some peak at G1, others at S, and others at G2/M, and only *CYCD3;3* and *CYCD5;1* follow the canonical expression pattern (Menges et al., 2005; Menges et al., 2006). Similarly, expression of the A-type *CYCA3* subfamily peaks at S, while *CYCA1* and *CYCA2* peak at G2/M (Menges et al., 2005). Only B-type cyclins and CDKB genes are consistently expressed at G2/M (Menges et al., 2005). This diversity of expression patterns could be a consequence of gene redundancy and cell cycle specialization. The question of how this system may have worked in early land plant ancestors is yet unresolved.

To understand the plant cell cycle regulation, it is crucial to employ diverse model species. Research in green algae like *Chlamydomonas reinhardtii* has provided critical insights into the fundamental architecture of the cell cycle (Cross and Umen, 2015; Tulin and Cross, 2015; Atkins and Cross, 2018; Breker et al., 2018; Cross, 2020), including conservation of the regulation of the anaphase-promoting complex by the plant specific CYCB/CDKB complex (Pecani et al., 2022). However, *Chlamydomonas* exhibit an uncommon multiple-fission cell cycle, that consists of a long G1 phase followed by successive rounds of S phases and M (Cross and Umen, 2015), making it hard to compare with other plant species. Non-vascular plants offer a promising opportunity to study the cell cycle in early divergent plants, with different cellular body plans. The moss *Physcomitrium patens* has revealed both conserved and unique aspects of plant cell cycle regulation (Schween et al., 2008; Ishikawa et al., 2011; Peramuna et al., 2023). However, the large repertoire of cyclins and CDKs in *Physcomitrium* creates challenges for understanding due to functional redundancy.

The model liverwort *Marchantia polymorpha*, features a reduced set of cell-cycle regulators (Bowman et al., 2017). Its simplicity positions Marchantia as a key organism to study the evolution of developmental and systems biology (Bowman et al., 2022; Romani et al., 2024) and could be instrumental to understand the evolution of the cell-cycle control system across land plants without the confounding effects of gene redundancy and sub-functionalization. However, functional studies in this species are lacking.

In this work, we made a comprehensive phylogenetic analysis of cell cycle genes across plants and other eukaryotes and observed that Marchantia retains a minimal set of regulators. Expression profiles and functional analyses allowed us to elucidate the organization of the core cell cycle regulation genes that operate in Marchantia and compare this system with its counterparts in other eukaryotes. Functional analyses of cyclin genes in the Marchantia vegetative gametophyte revealed that Mp*CYCD;1* is a rate-limiting factor for cell proliferation, promoting dedifferentiation, while Mp*CYCB;1* and Mp*CYCA* causes cell cycle arrest. On the other hand, MpCYCB;2 is a gene in the process pseudogenization and MpCYCD;1 is not a canonical D-type cyclin, without any significant impact in plant growth. Overall, these results provide a model of the streamlined cell-cycle control system in Marchantia and will be useful to navigate study of the cell cycle and future approaches to engineering plant growth and development.

## RESULTS

### Phylogenetic analysis of core cell cycle genes

To investigate the core regulation of cell cycle regulation in land plants, we conducted a phylogenetic analysis of cell cycle genes in a broad range of 27 eukaryotic species, including yeast, animals, brown and red algae, charophytes, bryophytes, lycophytes, ferns, and flowering plants (Figure 1). We focused on gene families that have been described as fundamental components of the cell-cycle control machinery, including: Group I cyclins, CDKs, CYCLIN-DEPENDENT KINASES REGULATORY SUBUNIT (CKS), RETINOBLASTOMA-RELATED (RBR) proteins, and the associated transcription factors (TFs) from the E2F, DP, DEL, and 3R-MYB gene families.

**Figure 1.**
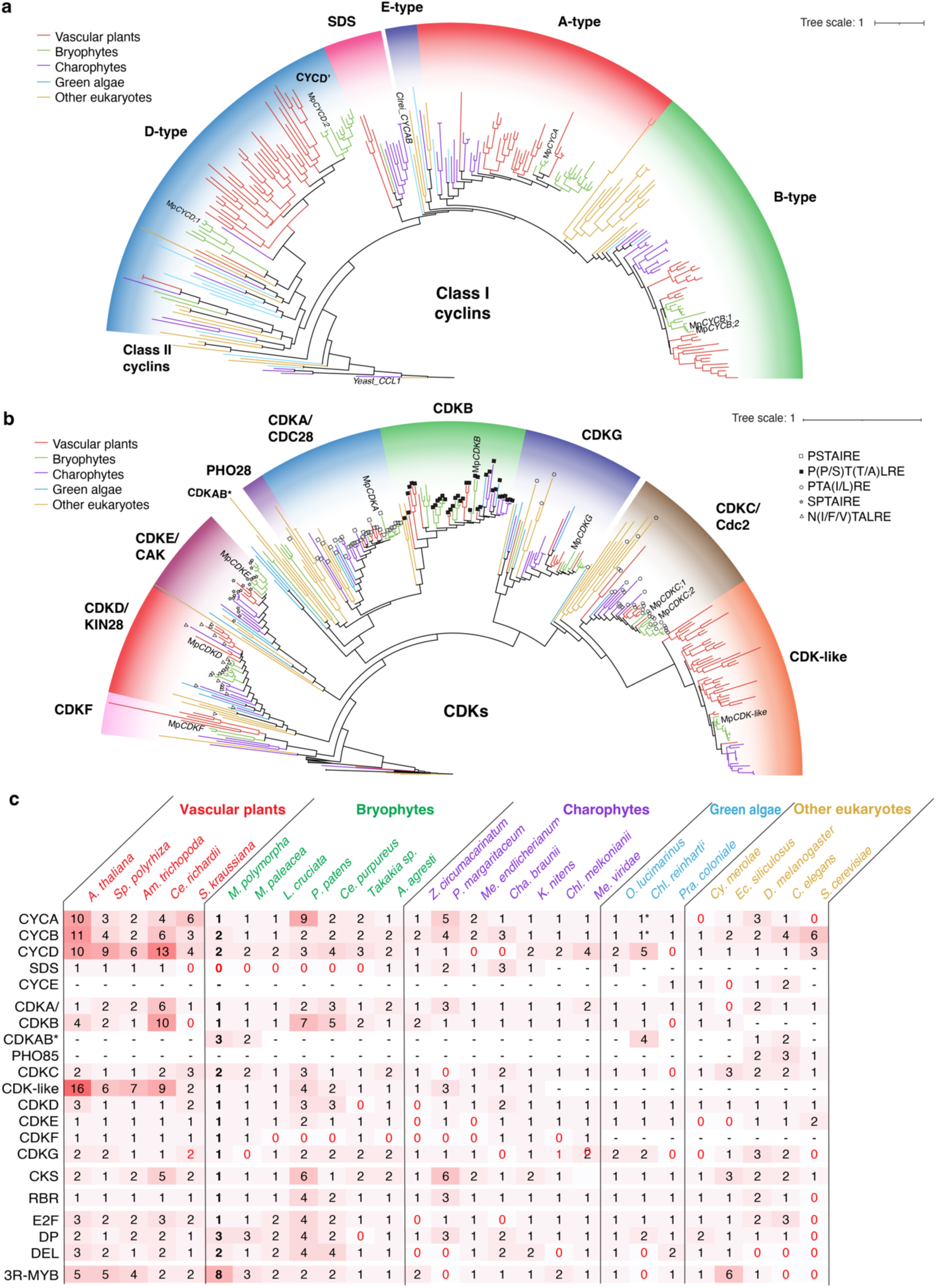
Phylogenetic analysis of core cell cycle genes across eukaryotes. a-b) Maximum likelihood Phylogenetic analysis of cyclins across representative plant and eukaryotic species. (a) and CDKs (b). In each diagram, subfamilies are indicated with different shaded background colours. Branches are coloured according to species in each clade as shown in the figure legends. Conserved signature motifs across CDKs are indicated in each node as shown in the figure key on the right. Full phylogenies and details are available at Suppl. File 1. c) Summary of the number of cell cycle genes across eukaryotic species. Background colour indicates the total number of genes in darker red. Hyphens indicate absence, while red zeros indicate potential losses. * Chlamydomonas reinhardtii has an extra divergent CYCAB1 (Cre08.g370401).

A-, B-, and D-type cyclins are well conserved across eukaryotes, except fungi, which only display two classes of cyclins (Ma et al., 2013) corresponding to the B- and D-type (Figure 1a,c). E-type cyclins are specific to animals, with potential presence in other green algal lineages but absent in land plants (Figure 1a-b). SOLO DANCERS (SDS) cyclins are a group of plant-specific cyclins associated with meiosis (Azumi et al., 2002) that likely originated in charophytic algae but those genes were subsequently lost in some non-seed plants (Figure 1c). Interestingly, there is a gene duplication of the D-type cyclins that appear to have given rise to a new clade of bryophyte-specific cyclins that we named CYCD’ (Figure 1a,c).

CDKAs and CDKBs are the main partners that interact with cyclins to help govern cell-cycle progression in plants. CDKA, homologous to yeast CDC28, is conserved across eukaryotes with the hallmark PSTAIRE motif (Figure 1b-c, Suppl. Figure 2). The G2/M specific CDKB is ubiquitously present in all green plants (Figure 1b-c). In non-flowering plants, they are associated with the PSTALRE motif (Figure 1b) that is well conserved across the family, as opposed to the P(P/S)T(A/T)LRE in flowering plants (Joubes et al., 2000). The only case in which this motif is found in CDKBs outside the green algae are the brown algae, *Ectocarpus siliculosus*, which contain two divergent CDKB-like CDKs with the PSTALRE motif (Figure 1b).

Other CDKs can act as activators within the cell cycle phosphorylation cascade (Joubes et al., 2000). CDKC (homologous to yeast CTK), CDKD (KIN28), CDKE (CAK1/SSN3), and CDKG are conserved across eukaryotes, with some lineage-specific losses (Figure 1b-c). Notably, the CDKC subfamily has expanded in plants, forming a distinct streptophyte-specific CDK-like clade, whose function is largely unknown. (Joubes et al., 2000). Among the remaining clades, CDKF is found in all streptophytes, and the PHO85 clade, which belong to the CDKA/CDC28 super-family, is conserved in yeast and animals but not found in most other eukaryotes (Liu and Kipreos, 2000). Divergent CDKAB members form a paraphyletic group in scattered species, lacking conserved motifs. CKS genes are broadly conserved, except in the charophyte algae *Mesostigma viridae* (Figure 1c).

Downstream of CDK signalling, the RBR pathway is essential for cell cycle progression and is also highly conserved, with a single RBR homologue in most species. There are three clades of RBR-related TFs that form part of the DREAM complex during the G1/S transition (E2F, DP, and DEL). The three clades share a common evolutionary origin (Rauber et al., 2016). E2F and DP TFs form highly conserved heterodimers to regulate the onset of DNA replication. Both protein families are present in animals and plants, suggesting an early origin of the interaction (Figure 1c). In contrast, DEL proteins, which are present in both brown algae and plants, have been lost in several plant lineages.

The 3R-MYB TFs (or c-Myb in animals), which regulate the G2/M transition (Ito et al., 2001), show a common origin before the divergence of plants and animals (Kobayashi et al., 2015; Feng et al., 2017), with some lineages losing one of the MYB domains. All plant species contain at least one 3R-MYB gene, except for the algae species *Chlamydomonas* and *Penium margaritaceum* (Figure 1c).

In charophytes, including the algal sister lineage of land plants (Feng et al., 2024), and bryophytes, most cell-cycle genes exist as single copies, suggesting that the common ancestor of land plants possessed a minimal set of these core components. This set has expanded considerably in some lineages, particularly in flowering plants, consistent with the pattern of whole-genome duplications. Remarkably, the liverwort *Marchantia polymorpha* retains a simplified set of cell cycle regulators, with highly reduced redundancy compared to vascular plants. It has single copies of most cell cycle regulators, except for CYCD (Mp*CYCD;1* and Mp*CYCD;2*) CYCB (Mp*CYCB;1* and Mp*CYCB;2*), and CDKC (MpCDKC;1 and MpCDKC;2), and a considerable repertoire of TFs, including 3 DP (Mp*DP1*, Mp*DP2*, and Mp*DP3*), 2 DEL (Mp*DEL1* and Mp*DEL2*), and 9 3R-MYBs (Mp*3R-MYB1* to Mp*3R-MYB9*).

### Expression profile of cell-cycle genes in Marchantia

To investigate the dynamic expression of key cell-cycle regulators in Marchantia, we re-analysed RNA-seq time-series data from two natural biological processes that feature a semi-synchronized re-entry to the cell cycle: meristem regeneration (Ishida et al., 2022) and sporeling germination (Bowman et al., 2017). During regeneration, cells begin proliferating approximately 16 hours after meristem excision (Nishihama et al., 2015). Similarly, dormant spores start dividing between 29 and 48 hours after exposure to light (Attrill et al., 2024).

During sporeling germination Mp*CYCD;1* transcripts accumulated after 24 hours of light exposure, before the first cell division, followed by peaks in Mp*CYCB;1*, Mp*CYCA*, Mp*CDKB*, Mp*CKS* at 48 hours (Figure 2a). We also employed *HISTONE H3.1* (Mp*H3.1)* transcript accumulation as a reference marker for DNA replication, which also peaked 48 hours after light exposure. A similar pattern was observed during regeneration (Figure 2b), consistent with previous reports (Nishihama et al., 2015). These findings are consistent with a role for Mp*CYCD;1* in G1, driving cell cycle re-entry, and highlight other genes that are induced during cell cycle progression. On the other hand, Mp*CDKA*, other CDKs, Mp*RBR,* and Mp*CYCD;2* remained constitutively expressed (Figure 2, Suppl. Figure 1, Suppl. Figure 2). Mp*E2F* and Mp*DP1* transcription factors showed moderate but detectable upregulation (Figure 1a-b).

**Figure 2.**
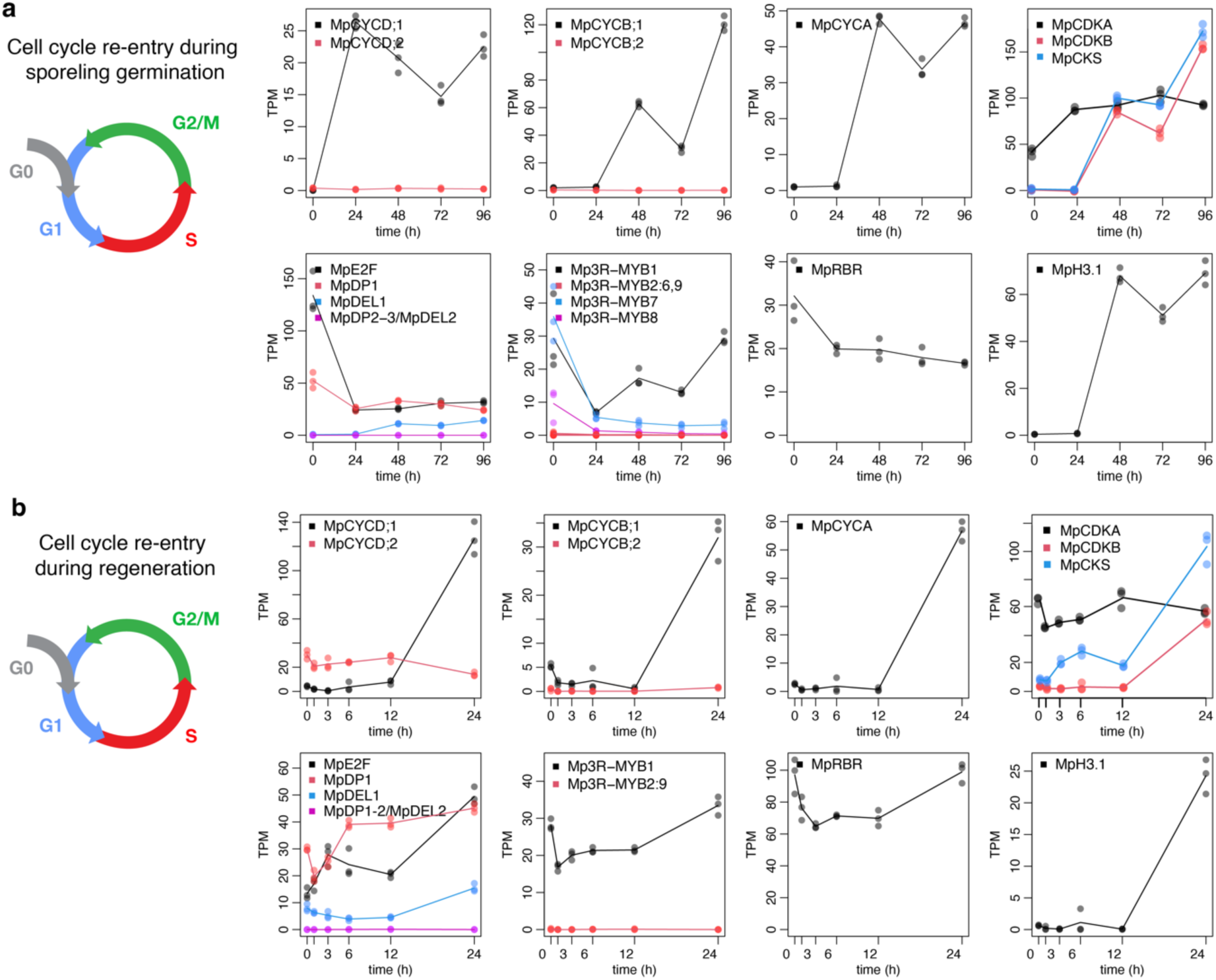
Expression of core cell cycle genes in Marchantia during re-entry. a-b) RNA-seq analysis of representative cell cycle genes during the time course of sporeling germination (a) and regeneration (b). Genes are grouped per families and distinguished by colour (see legend). Individual points represent biological replicates and lines averages.

Some predicted cell-cycle genes showed low or no detectable expression during both regeneration and sporeling germination, including: Mp*CYCB;2, MpCDKAB;1-3,* and Mp*DP2 MpDP3* (Figure 1, Suppl. Figure 1). Mp*3R-MYB1* remained the most abundantly expressed *3R-MYB* ortholog in the vegetative gametophyte. By contrast, Mp*3R-MYB3* to Mp*3R-MYB9* were specifically expressed during early sporophyte development, and Mp*3R-MYB7* and Mp*3R-MYB8* were detected at the spore stage too (Suppl. Figure 1). These 3R-MYB homologues belong to a different clade compared to 3R-MYB1 expressed in the gametophyte (Suppl. Figure 1). Something analogous was observed with Mp*DEL2*, which is expressed in the antheridium (Suppl. Figure 1). These data suggest that in the vegetative gametophyte, a simplified cell cycle regulation network is active, with only one copy of each TF. While in the sporophyte and during gametogenesis, some specialization could be expected.

To further investigate the spatial distribution of cell divisions and growth, we focused on the early stages of Marchantia gemma development. The meristem undergoes a transition from an immature state to the fully mature state around 5-days post-germination (Romani et al., 2024). We performed 5-ethynyl-2’-deoxyuridine (EdU) staining to label actively dividing cells. As expected, EdU-positive cells were concentrated around the division and differentiation zones (DDCZ) and the stem-cell zone (SCZ), including the apical cell (Figure 3a-b). This is the main focal point for growth in Marchantia, with a very simple architecture. Cells near the apical notch proliferate and differentiate, forming specialized cells and air pore structures. Epidermal cells exit the cell cycle and elongate as they become distal to the apical notch (Figure 3a-b).

**Figure 3.**
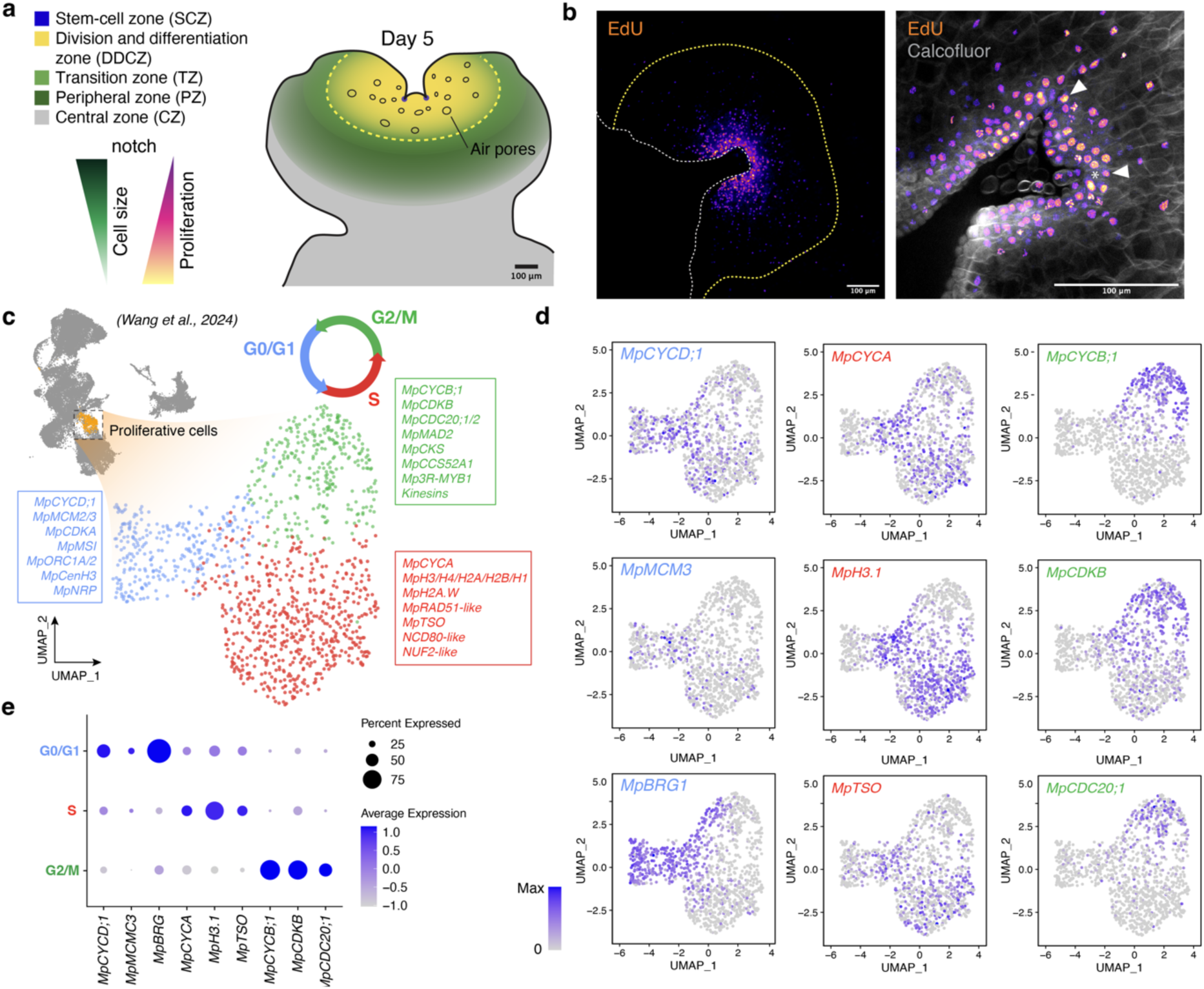
Gene expression fluctuations during cell proliferation in Marchantia. a) Diagram of cellular domains associated with cell proliferation in the Marchantia gemmaling b) 5-ethynyl-2’-deoxyuridine (EdU) staining of the Marchantia meristem in 5-days old gemmalings. The white dashed line delimitates the outline of the plant. The yellow dashed line represents the boundary of the mature epidermis. Scale bar is shown in each image. c-e) UMAP of scRNA-seq of Marchantia proliferative cells subcluster. The UMAP including all cells is shown in the top left corner for reference. Proliferative cells are highlighted in orange. e) UMAP of the proliferative cells and classification into 3 sub-clustering (G0/G1, light blue; S, red; and G2/M, green). Feature plots (d) and dot plot (e) of representative genes are shown in coloured square legends. Abbreviations: MCM (MINICHROMOSOME MAINTENANCE), CDC (CELL DIVISION CYCLE PROTEIN HOMOLOGUE), MSI (MUSASHI), ORC (ORIGIN RECONGNITION PROTEIN), CCS52A (CELL CYCLE SWITCH PROTEIN 52 A), NDC (NUCLEAR DIVISION CYCLE), NUF (KINETOCHORE PROTEIN NUF), BRG (BLEOMYCIN RESISTANT GENE), NRP (NAP-RELATED PROTEIN). TPM, transcripts per million; asinh, inverse hyperbolic sine.

To elucidate the expression of cell cycle regulators during gemmaling development, we generated transcriptional reporters using Golden-Gate compatible constructs (Sauret-Gueto et al., 2020; Romani et al., 2024). Transcriptional reporters showed that most key components were actively expressed in the EdU-positive zone (Suppl. Figure 3), corresponding to the DDCZ of the vegetative meristem. Consistent with the transcriptomic analysis, the promoter of Mp*CYCB;2* showed no detectable expression, suggesting it could be a pseudogene. *_pro_*Mp*CYCD;2* exhibited a more irregular pattern compared to *_pro_*Mp*CYCD;1*, which was consistently expressed in dividing cells. TFs linked to cell cycle regulation, including Mp*3R-MYB1*, and Mp*E2F,* followed expected expression patterns based on prior transcriptomic analyses in the gametophyte (Suppl. Figure 3). Expression of DP TFs reporters was not detectable at this stage (Romani et al., 2024). This is inconsistent with Mp*DP1* transcriptomics, likely due to the promoter being insufficient to capture all regulatory elements important for this gene.

### Phase specificity of cell-cycle genes

To gain a more refined understanding of the dynamic expression of cyclins during all phases of the cell-cycle, we leveraged single-cell RNA-seq data obtained from growing gemmalings. Previous work identified clusters of cells proliferating around the apical notch, including cell cycle related genes (Wang et al., 2023). Sub-clustering of this group revealed three clearly distinct expression profiles (Figure 3c). Using well-known markers, we labelled sub-clusters corresponding to G0/G1, S, and G2/M phases and identified a comprehensive set of additional phase specific markers (Suppl. Table 1). The G0/G1 sub-cluster was characterized by the expression of Mp*CYCD;1*, Mp*CDKA*, and Mp*MCM3*, homologues of *Arabidopsis* G1-phase markers (Zhang et al., 2021) and other genes associated G1 and initiation of replication (Figure 3c). The S-phase cluster was defined by replication-related genes, including Mp*CYCA*, several histones, and DNA damage repair genes such as Mp*RAD51*. The G2/M sub-cluster showed elevated expression of mitotic markers, including the single CDH1 unit of the Anaphase Promoting Complex/Cyclosome (APC/C) present in Marchantia (Mp*CCS52A1),* both *CDC;20* cofactors (Mp*CDC20;1* and Mp*CDC20;2*) (Figure 2e-f), and key core cell-cycle genes: Mp*CDKB,* Mp*CKS,* and Mp*3R-MYB1*. Among the other CDKs, Mp*CDKD* showed differential expression in the G2/M cluster and Mp*CDKG* and Mp*CDK-like* in G0/G1, similar to their homologues in *Arabidopsis* (Menges et al., 2005). Transcript levels of other key components of the cell cycle such as RBR and CKIs did not show preferred accumulation in specific phases of the cell-cycle.

Cyclins are not only subjected to tight transcriptional control during cell division, but also protein degradation. We made translational reporters for MpCYCD;1/2, MpCYCB;1, MpCYCA and MpCDKA to better capture the protein expression dynamics during cell division. These protein fusions correspond to the CDS and a C-terminal mVenus (mV) fluorescent protein. Compared to promoter fusions for the corresponding genes, the signals are more restricted to dividing cells. MpCYCD;1 is highly expressed in dividing cells as expected, and rhizoid precursors and air pores (Figure 4a, Suppl. Figure 4a). On the other hand, MpCYCD;2 translational reporter was undetectable in vegetative tissues. MpCYCB;1 has the typical “salt and pepper” (Figure 4a) as described in its homologues in flowering plants (Schnittger et al., 2002a). A similar pattern was observed for MpCYCA (Figure 4a). MpCDKA translational reporter is broadly expressed across the mature thallus and not restricted to dividing cells (Suppl. Figure 5a). Finally, MpCDKB is more restricted to dividing cells (Suppl. Figure 5b).

**Figure 4.**
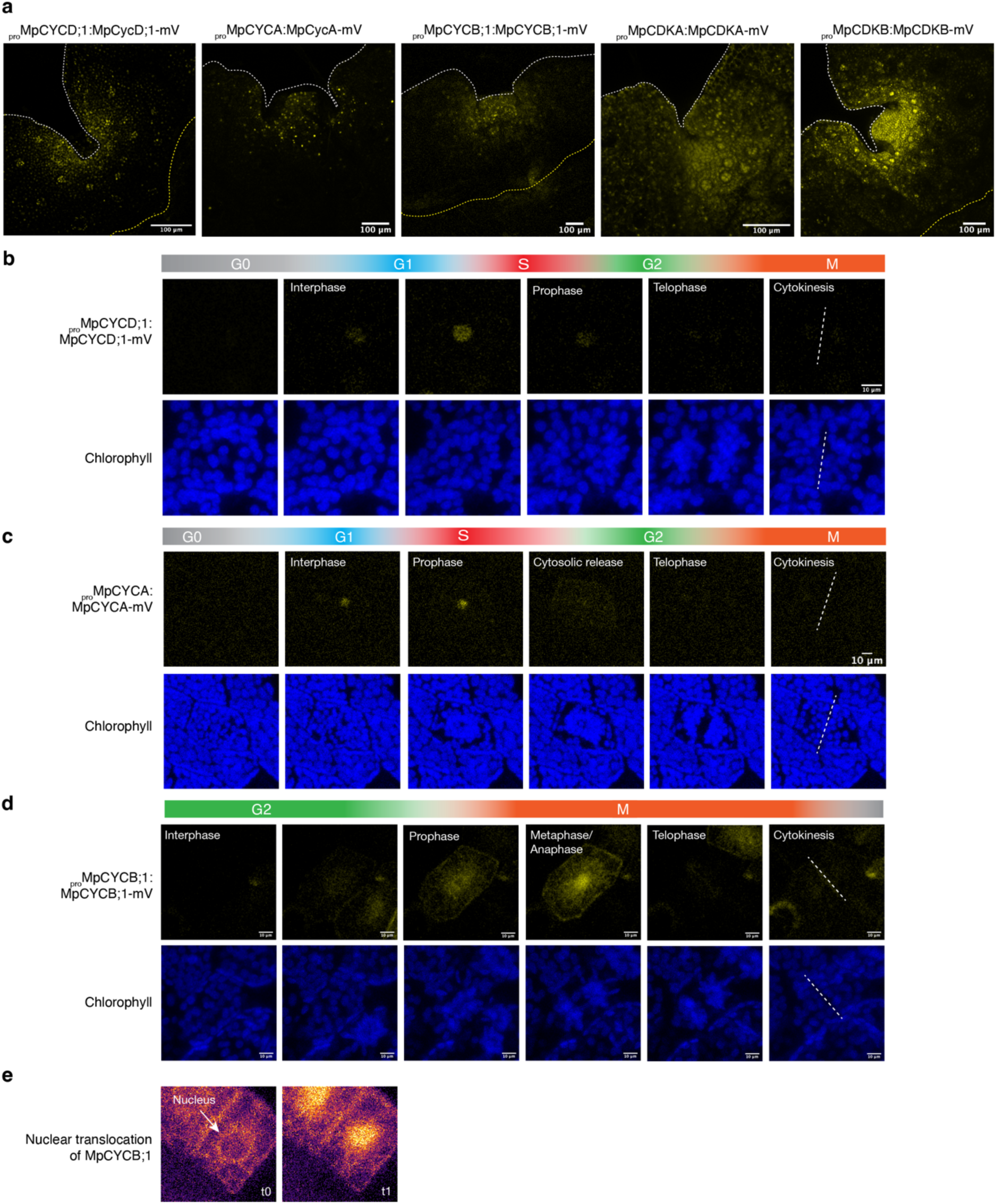
In vivo expression of cyclins and CDKs in Marchantia. Expression of cyclin and CDKs translational reporters (yellow) in 5-days old gemmalings. b-d) Time-lapse of MpCYCD;1 (b) MpCYCA (c), and MpCYCB;1 (d) translational reporters of an individual cell dividing during regeneration (yellow). Chlorophyll (blue) is shown to visualize the cell division phases using chloroplast movement. d) The translocation of MpCYCB;1 (inferno LUT) from the cytosol to the nuclei at two different frames. The full videos of the time-lapses are available at Suppl. Video 1-3. The white dashed lines delimit the outlines of the plants. The yellow dashed lines mark the boundary of the mature. Scale bars of the indicated sizes are shown in each image.

For time-lapse imaging, we took advantage of cell cycle re-entry occurring during regeneration, which allows a clear visualization of the cellular features associated with its expression dynamics during cell division using the translational reporters. Live imaging showed that MpCYCD;1 protein accumulated in the nuclei of cells before cell division and was rapidly degraded before the start of the prophase (Figure 4a, Suppl. Video 1). Similarly, MpCYCA also accumulated in the nuclei until the prophase and was rapidly released to the cytosol and degraded (Figure 4b, Suppl. Video 2). Finally, MpCYCB;1 was transiently expressed in the cytosol and accumulated during metaphase/anaphase, and signal was rapidly lost before cell plate formation (Figure 4c, Suppl. Video 3). Intriguingly, MpCYCB;1 was re-localized to the nuclei before loss of signal, likely due to degradation. The translocation of B-type cyclins during the metaphase-anaphase has been well described in animal cells (Clute and Pines, 1999) with the export of CYCB from the nucleus to the cytosol (Toyoshima et al., 1998). To our knowledge, this is the first report of this type of translocation in plant cells.

We also looked at translational reporters for MpCDKA and MpCDKB. Both were localized in the cytosol and nucleus and fluorescence was visible throughout the cell cycle. While MpCDKA strongly accumulated in the interphase (Suppl. Figure 6a, Suppl. Video 4), MpCDKB was found more prominent later in the prophase and during mitosis (Suppl. Figure 6b, Suppl. Video 5). Overall, these results align with the respective roles of the main cyclins and CDKS in Marchantia in the progression of the cell cycle. This also highlights the importance of protein degradation and nuclear export/import in controlling cyclin-dependent activity.

### Functional analysis of cyclins in Marchantia

We generated fluorescent protein fusions (CDS-mVenus) for overexpression of cyclins and CDKA. MpCYCD;1 is localized in the nucleus as most D-type cyclins in *Arabidopsis,* while MpCYCD;2 showed a broader distribution in the cytosol (Figure 5a). As shown before, CDKA is cytosolic (Figure 5a) as its orthologue in *Arabidopsis* (Boruc et al., 2010). Constitutive overexpression of Mp*CYCD;1* resulted in smaller plants with reduced cell size and severe developmental defects (Figure 5b). In contrast, overexpression of Mp*CYCD;2* or Mp*CDKA* had no noticeable effect on plant growth (Figure 5c).

**Figure 5.**
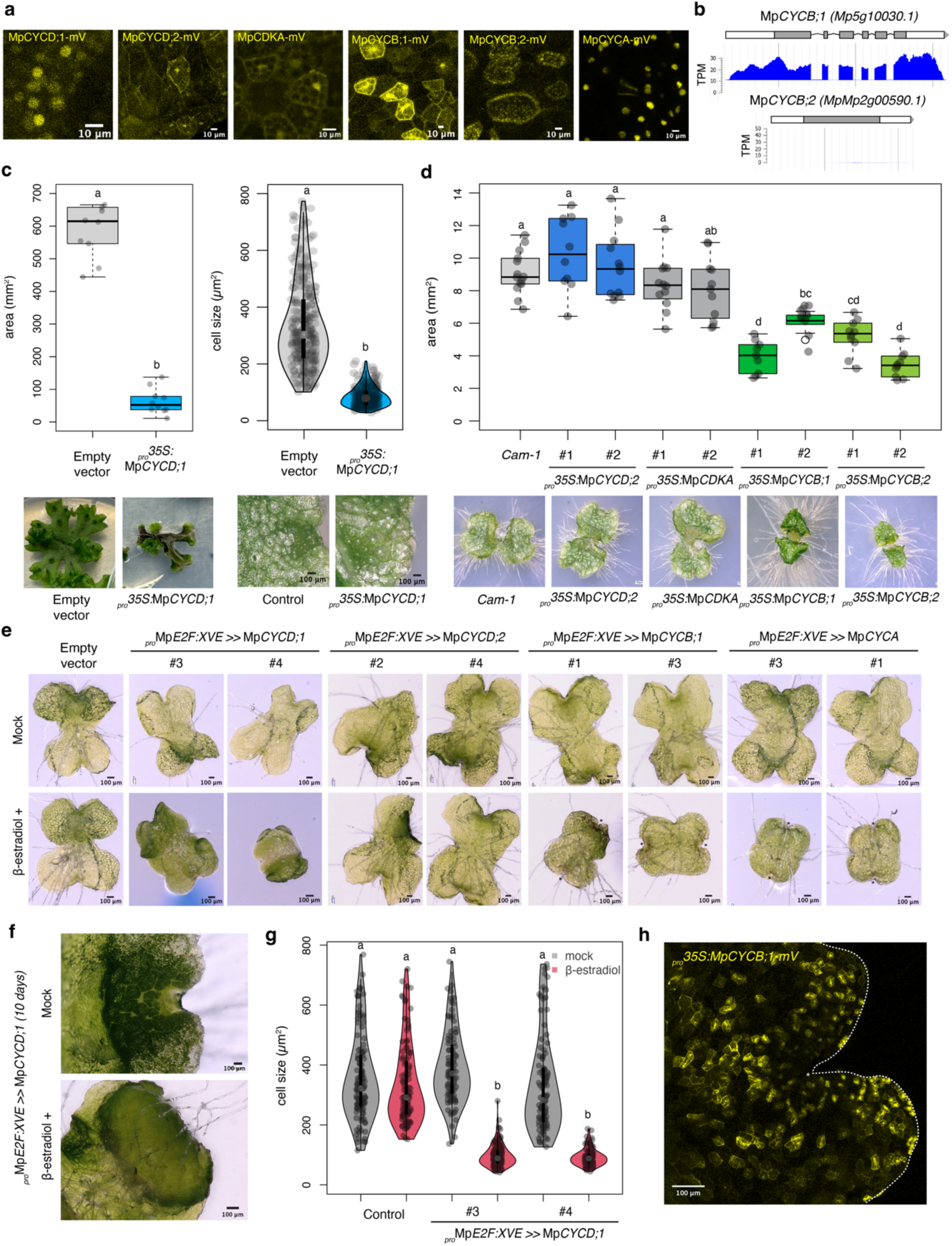
Functional characterization of cyclins and CDKA. a) Subcellular localization of selected cyclins and CDKA protein fusions with mVenus. b) Overview of the genomic locus of MpCYCB;1 and MpCYCB;2, including a diagram of intron structure and average RNA-seq coverage in mixed tissues. c) Boxplot of plant area (left) and cell size (right) of _pro_35S:MpCYCD;1-mVenus primary transformant plants. Representative pictures of the plants are shown. d) Boxplot of plant area of _pro_35S:MpCYCD;2-mVenus, _pro_35S:MpCDKA-mVenus, _pro_35S:MpCYCB;1-mVenus, and _pro_35S:MpCYCB;2-mVenus constitutive overexpression at 7 days. Representative pictures of the plants are shown at the bottom. e) Representative pictures of gemmalings of transgenic plants with inducible overexpression of cyclins grown for 3 days or 10 days (f) in 0.5x Gamborg B5 supplemented with 5 μM β-estradiol (EST+) or DMSO (mock). Transgenic constructs are indicated in the images. g) Violin plots of cell size of inducible MpCYCD;1 (_pro_MpE2F:XVE>>MpCYCD;1). in 3-days old plants. f) Confocal microscopy of _pro_35S:MpCYCB;1-mVenus (yellow) overexpression. The white dashed line delimitate the outline of the plant. The yellow dashed line represents the boundary of the mature epidermis. Control indicates a transgenic line transformed with an empty vector. Scale bar lengths are indicated in the pictures. Symbols above the bars indicate grouping by P-value < 0.001 in a Tukey’s Honest Significant Difference method.

Mp*CYCB;1* and Mp*CYCB;2* share high sequence similarity (90.1 %), but Mp*CYCB;2* lacks the conserved intron structure, and its endogenous expression is undetectable (Figure 2, Suppl. Figure 1, Figure 5b). Both B-type cyclins localize in the cytosol and nucleus (Figure 5a), consistent with its homologues in *Arabidopsis* (Boruc et al., 2010). Overexpression of either Mp*CYCB;1* or Mp*CYCB;2* impaired growth, resulting in smaller plants (Figure 5d). This indicates that despite signs of pseudogenization, Mp*CYCB;2* encodes for a functional protein. Imaging of plants overexpressing MpCYCB;1 revealed that cells with higher mVenus signal were arrested in mitosis at a metaphase-like stage (Figure 5h, Suppl. Video 6), unlike the transient expression observed native promoter control (Suppl. Video 7).

We also generated lines with inducible expression of Mp*CYCD;1*, Mp*CYCD;2*, Mp*CYCB;1* Mp*CYCA* using an β-estradiol XVE system under the control of _pro_MpE2F (Siligato et al., 2016; Ishida et al., 2022). Inducible overexpression of MpCYCD;1 further confirmed its role in controlling cell division, producing ectopic cell division and hyperplasia (Figure 5e,f), evidenced by significantly smaller cells under induction (Figure 5g) lacking proper differentiation. Under this setting, inducible overexpression of Mp*CYCD;2* did not cause any conspicuous developmental defect. On the other hand, inducible overexpression of Mp*CYCB;1* caused cell death. Inducible expression of Mp*CYCA* also caused a similar phenotype, suggesting that its accumulation is also arresting cells at mitosis (Figure 5g). This supports the key role of Mp*CYCB;1* and Mp*CYCA* in cell cycle control, while their degradation is required for the proper progression of cytokinesis.

Finally, to examine whether Mp*CYCD;2* play any physiological role for Marchantia vegetative development, we generated CRISPR-Cas9 knock-out lines. We obtain independent lines using two different gRNAs, generating a 20 bp (Mp*cycd;2-1^ge^*) or 1 bp (Mp*cycd;2-2^ge^*) deletions (Figure 6a, Supplemental Figure 7) that causes early stop codons. We did not find any significant defect in plant growth (Figure 6b) compared to wild-type plants. Considering overexpression results, this suggest that Mp*CYCD;2* does not affect cell proliferation and could be a pseudogene.

**Figure 6.**
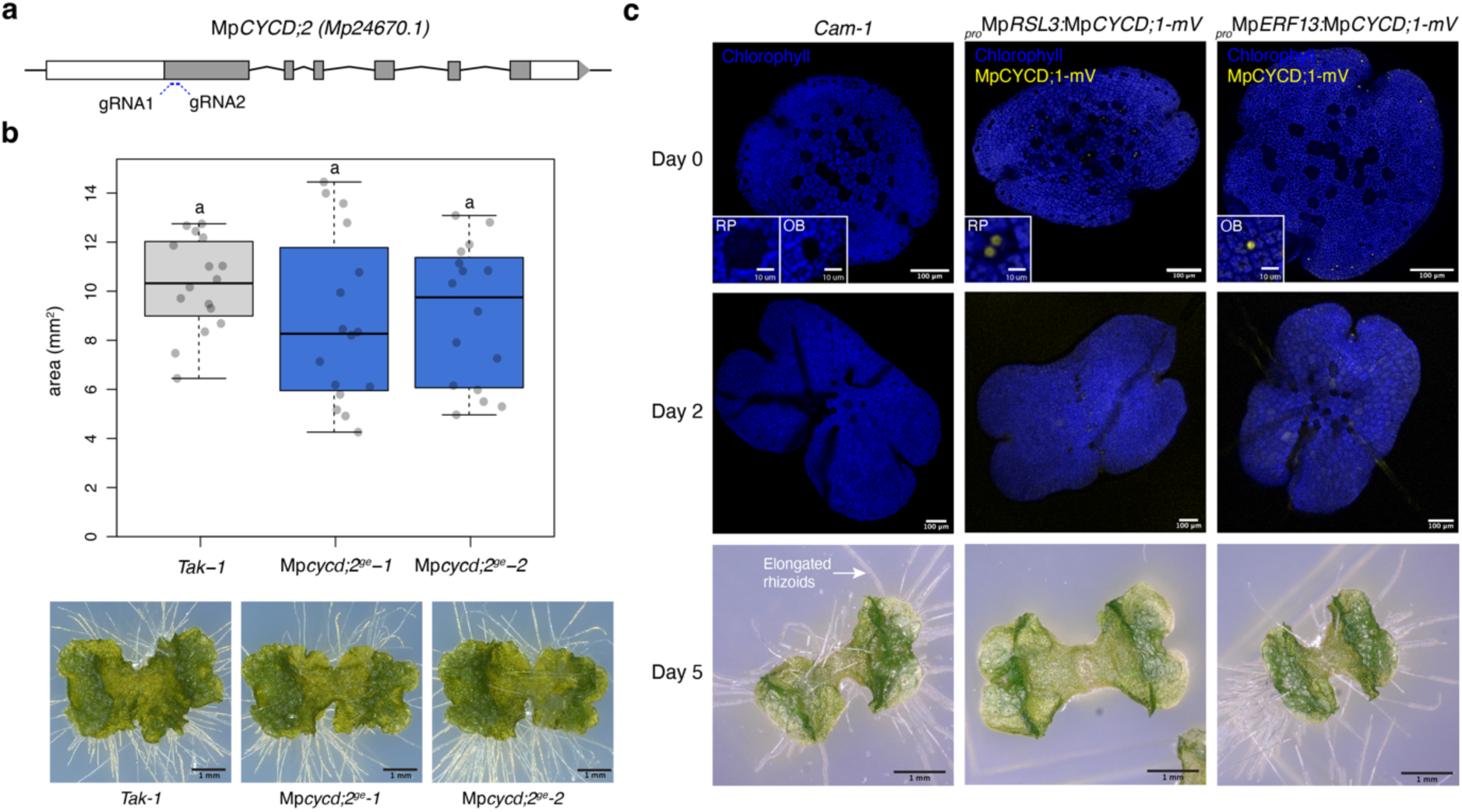
MpCYCD;2 knock-out and cell-type specific overexpression of MpCYCD;1 using cell-types specific promoters. a) Structure of MpCYCD;2 locus with the position of designed guide RNA (gRNA). Exons are shown as boxes. b) Boxplot of plant area of Mpcycd;2^ge^ and wild-type plants (Tak-1). Representative pictures of the plants are shown at the bottom. Scale bar lengths are indicated in the pictures. Symbols above the bars indicate grouping by P-value < 0.001 in a Tukey’s Honest Significant Difference method. c) Plants expressing _pro_MpRSL3:MpCYCD;1-mVenus (rhizoid precursor specific) and _pro_MpERF13:MpCYCD;1-mVenus (oil body specific). Wild-type Cam-1 was used as a control. Confocal full-stack images of representative individual gemmalings (0, and 3) are shown with mVenus (yellow) and chlorophyll (blue) channels merged. Representative picture of 5-days old gemmalings. Scale bar lengths are indicated in the pictures. RP: rhizoid precursors. OB: oil bodies.

### Cell Proliferation Interferes with Differentiation

Morphological defects observed after Mp*CYCD;1* overexpression showed that cell differentiation was disrupted when cell division was ectopically induced (Figure 5c,f). To explore this interaction, we employed cell-type-specific promoters to target differentiating cells, specifically rhizoid precursors and oil body cells (Sauret-Gueto et al., 2020; Romani et al., 2024). Both cell types, identifiable by their reduced chlorophyll content and distinct morphology, showed specific expression of Mp*CYCD;1* at the gemma stage. Misexpression of Mp*CYCD;1* under a rhizoid specific promoter (*_pro_MpRSL3*) caused ectopic cell divisions, resulting in the eventual loss of rhizoids (Figure 6c). Similarly, misexpression in oil body cells (*_pro_MpERF13*) triggered ectopic divisions, leaving smaller, scattered oil body cells in the gemma (Figure 6). Notably, these effects were not observed in Mp*CYCB;1* misexpression lines (Suppl. Figure 8).

During regeneration, both rhizoid precursors and oil body cells can undergo division and regenerate an entire plant after cell reprogramming (Nagai, 1919; Romani et al., 2024). De-differentiation of rhizoid precursors into epidermal cells is a normal phenomenon in some dorsal but not in ventral rhizoid precursors (Romani et al., 2024). The misexpression of Mp*CYCD;1* appears to extend this phenomenon, promoting widespread de-differentiation across ventral and dorsal precursors, impacting the emergence of rhizoids.

## DISCUSSION

We systematically characterised for the first time the functional integration of core cell-cycle genes in Marchantia. The genetic simplicity of Marchantia is second to none among plant model organisms, as its cell-cycle control system is neither expanded (as in most eukaryotes) nor reduced (as in yeast). On the other hand, displays the main features conserved among plants and has the potential to bridge the gap between unicellular algae and flowering plants at the systems level.

We showed that Marchantia displays simple set of cell-cycle genes (Figure 1). Particularly, the expression of cyclins follows an archetypal pattern with each gene peaking at the corresponding phase of its putative function, resembling the cell cycle regulation observed in animals and yeast. To achieve this, cyclin expression is tightly controlled at the transcriptional (Figure 2) and posttranscriptional (Figure 3) levels. In Marchantia this pattern is characterized by CYCD and CDKA during G1-phase, CYCA during S-phase, and CYCB and CDKB in G2/M (Figure 7a). Based on phylogenetic analysis and functional studies in Marchantia and other species, it can be inferred that this simple configuration represents the ancestral state of cell cycle regulation in eukaryotes, except for CDKB being specific to plants (Figure 7b).

**Figure 7.**
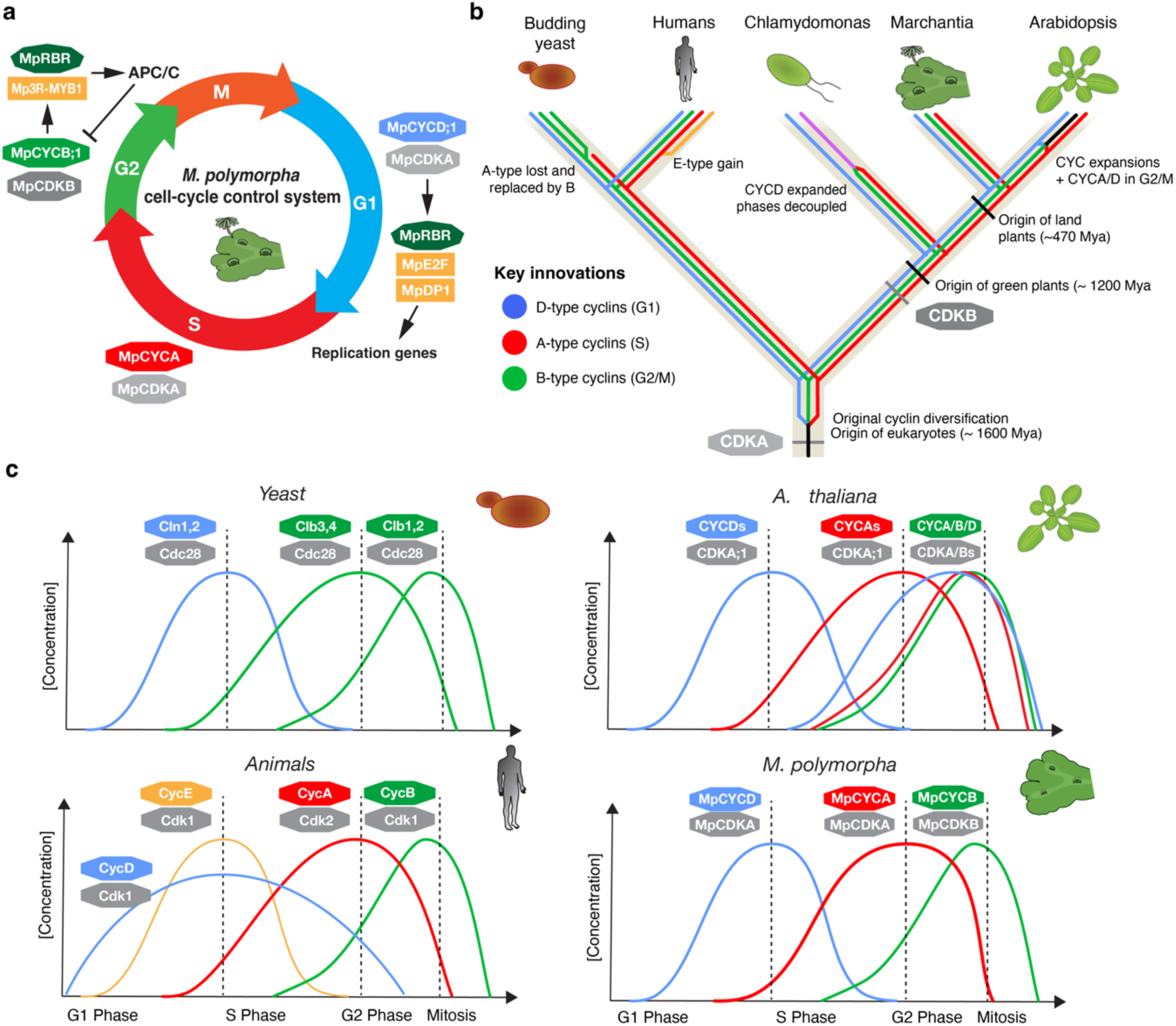
Core regulatory network of the cell cycle and phase specificity in Marchantia and other species. a) Diagram of cell cycle control system in Marchantia. b) Phylogram of the evolutionary trajectory of cell cycle genes in model eukaryotes (Saccharomyces cerevisiae, Homo sapiens, C. reinhartii, A. thaliana, and M. polymorpha) and its possible ancestral state. c) Comparison of expression patterns of cyclins and CDKs.

The streamlined system of cyclin-CDK components in Marchantia indicates that a complex regulatory network is not a prerequisite for multicellularity in plants but rather a derived feature in flowering plants. Thus, the extensive cyclin diversity observed in vascular plants likely arose through lineage-specific gene duplications, leading to redundancy and diversification. Subsequent neofunctionalization and subfunctionalization, may have facilitated cellular innovations and tissue specificity. In contrast, the set of key regulators that we described here for Marchantia are expressed in both the haploid and diploid generation (Suppl. Figure 1). This might imply that a very similar machinery controls cell division across both generations and meiosis, with few specific features.

The only two gene duplications among Class I cyclins in Marchantia, Mp*CYCB;2* and Mp*CYCD;2,* do not appear be relevant for cell cycle regulation. Mp*CYCB;2* is likely a pseudogene with negligible expression, while Mp*CYCD;2* belonging to a bryophyte-specific clade (CycD’). Overexpression, subcellular localization. and mutant phenotypes suggest that Mp*CYCD;2* is not a canonical D-type cyclin, and it is dispensable for vegetative growth.

Determining the truly essential components for the cell cycle progression remains a challenge across eukaryotes. While CDKA plays a central role across eukaryotes, it is not strictly essential in *Arabidopsis, Physcomitrium, and Chlamydomonas* (Nowack et al., 2012; Tulin and Cross, 2015; Bao et al., 2022). In Marchantia, CDKA is steadily expressed in dividing cells, with some preference for G1 (Figure 3), and does not appear to be a rate-limiting factor (Figure 5). CDKB, a G2/M-specific innovation conserved across green plants, is essential for spindle formation and nuclear division in *Chlamydomonas* (Tulin and Cross, 2014; Pecani et al., 2022). Our results suggest that MpCDKB is highly expressed at G2/M, likely interacting with MpCYCB;1. However, CDK protein levels are detected throughout the cell cycle and do not exhibit the strict phase-specific posttranscriptional regulation observed for cyclins.

D-type cyclins are conserved across eukaryotes and are undoubtedly important for plant growth and development, though their essentiality in plants remains unclear (Cross and Umen, 2015). Overexpression of Mp*CYCD;1* in Marchantia produces phenotypes similar to those observed for *CYCD3* homologs in flowering plants (Dewitte et al., 2003; Koroleva et al., 2004; Menges et al., 2006), reinforcing the idea that regulating cell cycle re-entry is the ancestral role of D-type cyclins in plants. This is important to understand the divergent roles of other D-type cyclins in flowering plants, and contrast with the role of D-type cyclins in animals (Datar et al., 2000).

We also demonstrated a close relationship between cyclin-mediated cell cycle re-entry and cellular differentiation in Marchantia (Figure 6). Our findings suggest that maintaining proper coordination of cell division rates is essential for preserving cell identity. This contrasts with Arabidopsis, where manipulating the cell cycle in specialized cells, such as trichomes, alters cell size and induces multicellularity but maintain cell identity (Schnittger et al., 2002b; Schnittger et al., 2002a). These differences may help explain Marchantia’s remarkable regenerative capacity and developmental plasticity (Nishihama et al., 2015).

Interestingly, overexpression of Mp*CYCA* or Mp*CYCB* is deleterious in Marchantia, leading to cell cycle arrest. In the case of A-type cyclins, they are usually associated with ploidy level and control of S-phase duration in plants (Imai et al., 2006). However, this might not be the case in Marchantia and other non-seed plants due to the absence of endoreduplication (Bainard et al., 2013; Nishihama et al., 2015). In *Chlamydomonas*, CYCA plays a critical role in cell division, but it is not essential (Atkins and Cross, 2018). It was also shown that overexpression of *CYCA3* in Arabidopsis cyclins affect meristem activity (Takahashi et al., 2010; Willems et al., 2020). Given that Arabidopsis *CYCA3* cyclins are also expressed at the S phase as MpCYCA in Marchantia. This suggest that controlling cell cycle progression during the S phase could be the ancestral role for A-type cyclins across plants. Subsequently, the role in ploidy could be restricted to the *CYCA2* clade and meiosis to *CYCA1* clade (d’Erfurth et al., 2010).

In the case of B-type cyclins, both function and phase specificity are well conserved across eukaryotes, playing a key role in regulating the G2/M transition, as we showed in Marchantia. Their essentiality for cell division was demonstrated in *Chlamydomonas* (Pecani et al., 2022). Historically, overexpression of *CYCB* is often associated with higher division rates (Doerner et al., 1996). However, in Marchantia, overexpression of Mp*CYCB;1* or Mp*CYCA* did not increase division rates but instead caused mitotic arrest. This arrest may result from the inability of some cells to degrade excessive cyclins, thereby preventing progression through mitosis (Figure 5h). The expression of non-degradable CYCBs in plants and other eukaryotes has also been shown to cause mitotic arrest, supporting that their degradation by the APC/C is critical (Figure 6). Similarly, A-type cyclins also need to be degraded by the APC/C but prior to the spindle assembly checkpoint in other eukaryotes (Geley et al., 2001). Our results suggest that *CYCA* degradation is also critical for cell-cycle progression in plants too (Figure 5).

It remains an open question whether A- or B-type cyclins are limiting factors for growth in Marchantia, as has been observed in other plants (Doerner et al., 1996). A possible explanation is that the balance between cell division or arrest is highly dose dependent. In addition to their role in mitotic control, CYCB1;1 induction in Arabidopsis has also been shown to cause mitotic arrest under DNA damage conditions (Schnittger and De Veylder, 2018). In animals, B-type cyclins are also not rate limiting for cell proliferation (Doonan and Hunt, 1996), suggesting that this could represent the ancestral state of CYCB protein function in eukaryotes. However, further studies are required to generalize the precise functions of A- and B-type cyclins in plants.

On the other hand, we showed that MpCYCD;1 is the single rate-limiting cyclin/CDK component for cell proliferation in Marchantia. Unlike MpCYCA and MpCYCB;1, its degradation may not be required for mitotic progression. Rather, its accumulation triggers re-entry into G1 and may be crucial for integrating developmental and environmental signals to regulate plant growth rate.

The role of the DREAM complex TFs (E2F/DP and 3R-MYBs) is also important to shape the downstream signalling networks of the cell cycle. The Marchantia genome encodes homologs of these regulators, but only one copy of each component is expressed in the gametophyte. The single-copy nature of E2F TFs in Marchantia and other land plants suggests that the complexity observed in Arabidopsis and animals arose independently (Sozzani et al., 2006; Rauber et al., 2016). The diversification of 3R-MYBs appears to be ancestral to land plants, and their expansion in Marchantia may be linked to sporophyte-specific programs (Supplementary File 1).

In conclusion, Marchantia provides a valuable model for studying cell cycle to demonstrate general principles of growth regulation that could be applied to complex organisms. Further studies using conditional knockouts (Nishihama et al., 2016) could clarify the roles of specific cell cycle genes in land plants. Additionally, targeted investigations into how environmental stress, DNA damage, hormones, or mechanical cues influence cell cycle progression will further enhance our understanding. The foundational work presented here establishes a toolkit of genes and promoters that can be leveraged for biotechnological applications. A deeper understanding of the cell cycle will enable more precise approaches to reprogramming plant growth and development.

## ACKNOWLEDGEMENTS

This work was funded by BBSRC BB/T007117/1 to J.H, BBSRC BB/F011458/1 for confocal microscopy, JSPS KAKENHI Grant Number JP22H02676 to Y.H., and Takeda Science Foundation (Life Science Research Grants, 2024034869) to Y.H. F.R. is a Leverhulme Early Career Fellow (ECF-2023-534) funded by the Leverhulme Trust and the Isaac Newton Trust (23.08(f)), I.B. is funded by the Herschel Smith Fund studentship. We thank Miguel A. Blazquez and Antonio Serrano Mislata (IBMCP, Valencia, Spain) for useful discussion and feedback on the manuscript. We thank Jia-Wei Wang and Long Wang (CAS, China) for advice and help on scRNA-seq data analysis.

## MATERIAL AND METHODS

### Phylogenetic analysis

*Ceratopteris richardii* v2.1 (Marchant et al., 2022), *Amborella trichopoda* v1.0 (Amborella Genome, 2013), *Chlamydomonas reinhardtii* v5.5 (Merchant et al., 2007), *Physcomitrium patens* v6.1 (Bi et al., 2024) and *Spirodela polyrhiza v2* (Wang et al., 2014) proteomes were obtained from Phytozome. *Phaeodactylum tricornutum* ASM15095 v2 (Bowler et al., 2008), *Prasinoderma coloniale* v1.1 (Li et al., 2020b), *Ostreococcus lucimarinus* ASM9206 v1 (Palenik et al., 2007), *Mesotaenium endlicherianum* v2 (Cheng et al., 2019), *Mesostigma viridea* (Liang et al., 2020)(ref), *Ectocarpus siliculosus (Cock et al., 2010)*, and *Chara braunii* v1.0 (Nishiyama et al., 2018) proteomes were obtained from Phycocosm. *Saccharomyces cerevisiae* R64.1.1 (Goffeau et al., 1996), *Caenorhabditis elegans (Consortium, 1998)*, and *Drosophila melanogaster* BDGP6.46 (Adams et al., 2000) proteomes were obtained from Ensembl, and *Cyanidioschyzon merolae* ASM9120 v1 (Matsuzaki et al., 2004) was obtained from Ensembl Plants. The proteomes from *Marchantia polymorpha Tak* v6.1 (Montgomery et al., 2020), *Selaginella kraussiana* v2 (Liu et al., 2023), *Anthoceros agresti* Oxford (Li et al., 2020a), *Klebsormidium nitens* NIES-2285 (Hori et al., 2014), *Zygnema circumcarinatum* (Feng et al., 2024), *Penium margaritaceum* (Jiao et al., 2020), *Marchantia paleacea* (Radhakrishnan et al., 2020), *Lunularia cruciata* v1 (Linde et al., 2020) from their respective sources.

Hidden Markov Models matrix for PFAMs including cyclins (PF00134.28), CKS (PF01111.24), E2F (PF02319.25), and RBR (PF01858.22) were used as query for searching using hmmsearch function from HMMER v3.4 (Finn et al., 2011). The cut-off E-value was then adjusted aiming to have 100% sensitivity for annotated genes and exclude low confidence findings: Cyclins < 1E-10, CKS < 1E-10, E2F < 1E-5, RBR < 1E-50. Using the hmmsearch output, Seqinr in R (Charif and Lobry, 2007) package was to retrieve the sequences from the proteomes.

3R-MYBs proteins were BLASTed using the human MYBB sequence as a query (threshold E-value < 2e-34) aiming to have 100% sensitivity for annotated 3R-MYB and include some R2R3-MYBs. Similarly, for CDK proteins, using yeast CDC28 and Arabidopsis CDKF as query (threshold E-value < 1e-44) aiming to have 100% sensitivity for annotated CDKs. Sequences were extracted from the proteomes using Biostrings package in R (Lifschitz et al., 2022).

Finally, JalView (Procter et al., 2021) was used to align sequences with the MAFFT software (Katoh et al., 2002) and trimming the alignment to include only columns with 50% coverage. Maximum-likelyhood phylogenetic analysis was performed using IQtree 2 (Minh et al., 2020) with default parameters. Trees were rooted and visualised using iTOL v6 (Letunic and Bork, 2024). Complete trees available at (Suppl. File 1).

To count the number of MYB repeats (SM00717) for 3R-MYB proteins, we used InterProScan (Quevillon et al., 2005). To identify signature motifs on CDKs we used a simple grepl search function. Marchantia genes names were annotated in MarpolBase.

### Analysis of public RNA-sequencing data

TPM values for sporeling germination and different developmental stages were extracted from Marpolbase Expression database (Kawamura et al., 2022) or downloaded from SRA for regeneration (DRR330148-DRR330173), mapped in *M. polymorpha* Tak-1 genome v5.1 using HISAT2 (Kim et al., 2019), ht-seq and EdgeR. Data was subsequently analysed with R to generate plots using customs scripts. Heatmap was generated using Marpolbase Expression database (Kawamura et al., 2022).

The scRNA-seq matrix for Marchantia gemmaling development (Wang et al., 2023) was extracted from Beijing Institute of Genomics Data Center (OMIX004749) and re-analysed using Seurat V5 package in R (Hao et al., 2024). Briefly, the dataset was subseted to only cluster 10 and analysed with the following functions and parameters: FindNeighbours (dims = 1:100, k.param = 200), FindClusters, RunUMAP (reduction = ‘pca’, metric = ‘correlation’, n.neighbours = 30L, dims = 1:100). Data was visualized using DimPlot, FeaturePlot and DotPlot functions. Markers in Suppl. Table S1 was obtained using FindAllMarkers function and genes annotated using MarpolBase.

### Plant material and growth conditions

*Marchantia polymorpha* subs*. rudelaris* accessions *Cam-1* (male) and *Cam-2* (female) were used for most of the experiments. Under normal conditions, plants were grown on solid 0.5× Gamborg B-5 basal medium (Phytotech #G398) at pH 5.8 with 1.2% (w/v) agar micropropagation grade (Phytotech #A296), under continuous LED light at 21 °C with light intensity of 150 *μ*mol/m^2^/s (Systion #SE-EGB). For spore production, plants were grown in Microbox micropropagation containers (SacO_2_) in long-day conditions (16 h light/8 h dark) under light supplemented with far-red light as described (Sauret-Gueto et al., 2020).

For Mp*cycd;2* knock-outs, the *M. polymorpha* subs*. rudelaris* male accession Takaragaike-1 (Tak-1) accession was used as wild type. Similarly, *M. polymorpha* plants were grown on half-strength Gamborg B5 medium (pH 5.5) solidified with 1.4% (w/v) agar at 22 ℃ under continuous light.

### Plasmid construction

To generate new L0 parts, CDS and promoter regions from genes were extracted from *M. polymorpha* Tak-1 genome version 5.1 (Bowman et al., 2017) genome and manually domesticated to remove internal BsaI and SapI sites using synonymous mutations for the CDS. The sequences of synthetic L0 parts used in this work are available in Suppl. Table S2. L0 parts were synthesized by GENEWIZ following the standard syntax for plant synthetic biology with CDS and PROM5 or PROM and 5UTR overhangs and cloned into the plasmid pUAP1 (Addgene #63674) (Patron et al., 2015) by homology recombination. Other parts were derived from previous works (Sauret-Gueto et al., 2020; Romani et al., 2024) as specified in Suppl. Table S2.

The acceptor pBy01 (Romani et al., 2024) was used to assemble using L0 corresponding to PROM5 or PROM and 5UTR parts and pBy10 (Tse et al., 2024) for entire cassettes containing a hygromycin resistant cassette for plant selection. These acceptors are binary vectors contain a LacZ selection cassette flanked by BsaI sites to clone final vectors in one-step using L0 parts following the standard syntaxis (Patron et al., 2015). Following the same logic, a new custom acceptor for chlorosulfuron selection was generated based on pBy10 and OP-62 (Sauret-Gueto et al., 2020; Romani et al., 2024). Another two custom acceptors were generated for cloning only CDS with the *_pro_35S* (pBy12) or estradiol inducible system using *_pro_MpE2F:XVE* driving the expression of *_pro_LexA,* similar to previous published Gateway acceptor pMpGWB168 *(Ishida et al., 2022)* and a *_3utr_NOS* terminator, containing also an hygromycin (pBy13) or chlorosulfuron (pBy23) resistant cassette for plant selection. The full-length of the final constructs were verified by sequencing using the Oxford Nanopore technology (Plasmidsaurus Inc.). The final plasmid map is provided in the supplementary data (Supplemental File 2-3).

Plasmids were assembled as detailed in Suppl. Table S2 using Type-IIS cloning as described previously for L3 plasmids (Romani et al., 2024). Briefly using a Master Mix containing 10% (v/v) 10× T4 DNA ligase buffer (NEB #M0202), 5% (v/v) T4 DNA ligase at 400 U/μL (NEB #M0202), 5% (v/v) BsaI at 20 U/μL (NEB #R3733), 10% (v/v) acceptor at 40 ng/μL, 20% (v/v) pre-mixed L0 parts (∼100 ng/μL) and water to a final volume of 5 μL. Cycling conditions were 26 cycles of 37 °C for 3 min and 16 °C for 4 min. Termination and enzyme denaturation: 50 °C for 5 min, and 80 °C for 10 min. Other L1 and L2 plasmids were cloned as described before (Sauret-Gueto et al., 2020). TOP10 chemically competent *Escherichia coli* cells were transformed using the assembly reaction and plated on LB-agar plates containing the corresponding antibiotic and 40 μg/mL of 5-bromo-4-chloro-3-indolyl β-D-galactopyranoside (X-Gal). The presence of the correct insert was confirmed by restriction XhoI digestion (Thermo Scientific #FD0694) and Sanger sequencing, with primers available at Supplemental Table S2.

For genome editing of Mp*CYCD;2* (*Mp1g24670*), guide RNA was designed at the coding sequence of the first intron using CRISPRdirect (Naito et al., 2015) and primers at Supplemental Table S2. The plasmids were constructed according to Sugano et al., using pMpGE_En03 and pMpGE010 (Sugano et al., 2018).

### Plant transformation

*Agrobacterium tumefaciens* (GV3101) cells were transformed using the freeze-thaw method and used for plant transformation of *Cam-1/2* spores as described in Annese et al., (2025). Plants were screened for positive fluorescence and at least two independent lines were selected. For expression markers, representative transgenic lines are shown, showing the consensus expression pattern out of 4-5 independent lines.

For genome editing of Mp*CYCD;2*, Tak-1 plants were transformed using the cutting thallus transformation protocol (Kubota et al., 2013) and genotyped using primers at Supplemental Table S2.

### Plant phenotyping

Representative images of Marchantia plants were taken using a Keyence VHX-5000 digital microscope equipped with a 20x-200x Ultra-Small, High-Performance Zoom Lens (VH-Z20R/Z20T) or a Leica DMS1000 digital microscope. Thallus area was quantified from pictures (normally 7-days old gemmalings) with ImageJ software with at least 10 biological replicates. Cell sizes were quantified from manual segmentation of cells in 200x pictures using ImageJ. Only cells surrounding the DDCZ but excluding smaller cells closer to the apical meristem. At least 50 cells were measured for each biological replicate in at least 3 biological replicates.

For β-estradiol (#E8875, Merck) treatments, a 50 mM stock was dissolved in DMSO and added to melted agar-Gamborg B-5 media at a final concentration of 5 μM as described in Ishida et al., (2021). Plants were grown in microscopy contact plates and visualized after 3 or 10 days in normal growth conditions.

### Laser scanning confocal microscopy

Confocal images of Marchantia were acquired on a Leica SP8X spectral confocal microscope upright system equipped with a 460 to 670 nm super continuum white light laser (80% laser power), two CW laser lines 405, and 442 nm, and 5 Channel Spectral Scanhead (four hybrid detectors and one photomultiplier). For slides, imaging was conducted using either a 10× air objective (HC PL APO 10×/0.40 CS2) or a 20× air objective (HC PL APO 20×/0.75 CS2). When observing fluorescent protein with overlapping emission spectra, sequential scanning mode was selected. Excitation laser wavelength and captured emitted fluorescence wavelength window were: for mVenus (514 nm, 527 to 552 nm), and for chlorophyll autofluorescence (633, 687 to 739 nm).

### Time-course and time-lapses

When imaging time-courses, plants grown under normal culture conditions in small petri dishes, removed the lid for imaging, and returned the plants to the growth chamber and imaged as described above. For live imaging, six stacked Gene Frames (#AB0578, ThermoFisher) were placed on a glass slide and filled halfway with molten Gamborg B-5 agar medium. Plants were then placed on the solidified agar surface and meristems were removed using a Laser Microdissection Leica LMD7000. Samples were mounted in perfluorodecalin (Littlejohn et al., 2010) with a glass coverslip on top. The slides were then continuously imaged on the Leica SP8X confocal microscope for 1 to 4 d.

### EdU staining

5-days old gemmalings were incubated in liquid half-strength Gamborg’s B-5 medium under continuous light for 3 h with 20 μM 5-ethynyl-2’-deoxyuridine (EdU) from the Click-iT EdU Imaging Kit with Alexa Fluor 488 (#C10337, Invitrogen). Then they were fixed with 4% formaldehyde for 1 h, washed twice with phosphate buffer saline (PBS) and 0.5% Triton X-100 in PBS for 20 min. After washes, samples were incubated with a freshly prepared reaction mixture following the manufacturer instructions but without the Hoechst 33342 component. After staining, samples were protected from light, washed twice with PBS and soaked in iTomei-D (#T3940, Tokyo Chemical Industry) clearing solution and mounted in 70% w/v iohexol (#I0903, Tokyo Chemical Industry) in PBS as described before (Sakamoto et al., 2022). Samples were covered with a glass coverslip and imaged on the Leica SP8X confocal microscope as described before.

### Statistics

To obtain plots and statistical analysis, data was processed using R xv 4.4.1 software. For average and boxplots, the stats package was used with default parameters. For violinplots, the vioplot package was implemented also with default parameters. Statistical significance was calculated using ANOVA using the stats package Tukey ‘Honest Significant Difference’ method (alpha = 0.05) for levels calculations.

## REFERENCES

Adams, M.D., Celniker, S.E., Holt, R.A., Evans, C.A., Gocayne, J.D., Amanatides, P.G., Scherer, S.E., Li, P.W., Hoskins, R.A., Galle, R.F., George, R.A., Lewis, S.E., Richards, S., Ashburner, M., Henderson, S.N., Sutton, G.G., Wortman, J.R., Yandell, M.D., Zhang, Q., Chen, L.X., Brandon, R.C., Rogers, Y.H., Blazej, R.G., Champe, M., Pfeiffer, B.D., Wan, K.H., Doyle, C., Baxter, E.G., Helt, G., Nelson, C.R., Gabor, G.L., Abril, J.F., Agbayani, A., An, H.J., Andrews-Pfannkoch, C., Baldwin, D., Ballew, R.M., Basu, A., Baxendale, J., Bayraktaroglu, L., Beasley, E.M., Beeson, K.Y., Benos, P.V., Berman, B.P., Bhandari, D., Bolshakov, S., Borkova, D., Botchan, M.R., Bouck, J., Brokstein, P., Brottier, P., Burtis, K.C., Busam, D.A., Butler, H., Cadieu, E., Center, A., Chandra, I., Cherry, J.M., Cawley, S., Dahlke, C., Davenport, L.B., Davies, P., de Pablos, B., Delcher, A., Deng, Z., Mays, A.D., Dew, I., Dietz, S.M., Dodson, K., Doup, L.E., Downes, M., Dugan-Rocha, S., Dunkov, B.C., Dunn, P., Durbin, K.J., Evangelista, C.C., Ferraz, C., Ferriera, S., Fleischmann, W., Fosler, C., Gabrielian, A.E., Garg, N.S., Gelbart, W.M., Glasser, K., Glodek, A., Gong, F., Gorrell, J.H., Gu, Z., Guan, P., Harris, M., Harris, N.L., Harvey, D., Heiman, T.J., Hernandez, J.R., Houck, J., Hostin, D., Houston, K.A., Howland, T.J., Wei, M.H., Ibegwam, C., Jalali, M., Kalush, F., Karpen, G.H., Ke, Z., Kennison, J.A., Ketchum, K.A., Kimmel, B.E., Kodira, C.D., Kraft, C., Kravitz, S., Kulp, D., Lai, Z., Lasko, P., Lei, Y., Levitsky, A.A., Li, J., Li, Z., Liang, Y., Lin, X., Liu, X., Mattei, B., McIntosh, T.C., McLeod, M.P., McPherson, D., Merkulov, G., Milshina, N.V., Mobarry, C., Morris, J., Moshrefi, A., Mount, S.M., Moy, M., Murphy, B., Murphy, L., Muzny, D.M., Nelson, D.L., Nelson, D.R., Nelson, K.A., Nixon, K., Nusskern, D.R., Pacleb, J.M., Palazzolo, M., Pittman, G.S., Pan, S., Pollard, J., Puri, V., Reese, M.G., Reinert, K., Remington, K., Saunders, R.D., Scheeler, F., Shen, H., Shue, B.C., Siden-Kiamos, I., Simpson, M., Skupski, M.P., Smith, T., Spier, E., Spradling, A.C., Stapleton, M., Strong, R., Sun, E., Svirskas, R., Tector, C., Turner, R., Venter, E., Wang, A.H., Wang, X., Wang, Z.Y., Wassarman, D.A., Weinstock, G.M., Weissenbach, J., Williams, S.M., Woodage T, Worley, K.C., Wu, D., Yang, S., Yao, Q.A., Ye, J., Yeh, R.F., Zaveri, J.S., Zhan, M., Zhang, G., Zhao, Q., Zheng, L., Zheng, X.H., Zhong, F.N., Zhong, W., Zhou, X., Zhu, S., Zhu, X., Smith, H.O., Gibbs, R.A., Myers, E.W., Rubin, G.M., and Venter, J.C. (2000). The genome sequence of Drosophila melanogaster. Science 287, 2185–2195.

Amborella Genome, P. (2013). The Amborella genome and the evolution of flowering plants. Science 342, 1241089.

Atkins, K.C., and Cross, F.R. (2018). Interregulation of CDKA/CDK1 and the Plant-Specific Cyclin-Dependent Kinase CDKB in Control of the Chlamydomonas Cell Cycle. Plant Cell 30, 429–446.

Attrill, S.T., Mulvey, H., Champion, C., and Dolan, L. (2024). Microtubules and actin filaments direct nuclear movement during the polarisation of Marchantia spore cells. Development 151.

Azumi, Y., Liu, D., Zhao, D., Li, W., Wang, G., Hu, Y., and Ma, H. (2002). Homolog interaction during meiotic prophase I in Arabidopsis requires the SOLO DANCERS gene encoding a novel cyclin-like protein. EMBO J 21, 3081–3095.

Bainard, J.D., Forrest, L.L., Goffinet, B., and Newmaster, S.G. (2013). Nuclear DNA content variation and evolution in liverworts. Mol Phylogenet Evol 68, 619–627.

Bao, L., Inoue, N., Ishikawa, M., Gotoh, E., Teh, O.K., Higa, T., Morimoto, T., Ginanjar, E.F., Harashima, H., Noda, N., Watahiki, M., Hiwatashi, Y., Sekine, M., Hasebe, M., Wada, M., and Fujita, T. (2022). A PSTAIRE-type cyclin-dependent kinase controls light responses in land plants. Sci Adv 8, eabk2116.

Bi, G., Zhao, S., Yao, J., Wang, H., Zhao, M., Sun, Y., Hou, X., Haas, F.B., Varshney, D., Prigge, M., Rensing, S.A., Jiao, Y., Ma, Y., Yan, J., and Dai, J. (2024). Near telomere-to-telomere genome of the model plant Physcomitrium patens. Nat Plants 10, 327–343.

Boruc, J., Mylle, E., Duda, M., De Clercq, R., Rombauts, S., Geelen, D., Hilson, P., Inze, D., Van Damme, D., and Russinova, E. (2010). Systematic localization of the Arabidopsis core cell cycle proteins reveals novel cell division complexes. Plant Physiol 152, 553–565.

Bowler, C., Allen, A.E., Badger, J.H., Grimwood, J., Jabbari, K., Kuo, A., Maheswari, U., Martens, C., Maumus, F., Otillar, R.P., Rayko, E., Salamov, A., Vandepoele, K., Beszteri, B., Gruber, A., Heijde, M., Katinka, M., Mock, T., Valentin, K., Verret, F., Berges, J.A., Brownlee, C., Cadoret, J.P., Chiovitti, A., Choi, C.J., Coesel, S., De Martino, A., Detter, J.C., Durkin, C., Falciatore, A., Fournet, J., Haruta, M., Huysman, M.J., Jenkins, B.D., Jiroutova, K., Jorgensen, R.E., Joubert, Y., Kaplan, A., Kroger, N., Kroth, P.G., La Roche, J., Lindquist, E., Lommer, M., Martin-Jezequel, V., Lopez, P.J., Lucas, S., Mangogna, M., McGinnis, K., Medlin, L.K., Montsant, A., Oudot-Le Secq, M.P., Napoli, C., Obornik, M., Parker, M.S., Petit, J.L., Porcel, B.M., Poulsen, N., Robison, M., Rychlewski, L., Rynearson, T.A., Schmutz, J., Shapiro, H., Siaut, M., Stanley, M., Sussman, M.R., Taylor, A.R., Vardi, A., von Dassow, P., Vyverman, W., Willis, A., Wyrwicz, L.S., Rokhsar, D.S., Weissenbach, J., Armbrust, E.V., Green, B.R., Van de Peer, Y., and Grigoriev, I.V. (2008). The Phaeodactylum genome reveals the evolutionary history of diatom genomes. Nature 456, 239–244.

Bowman, J.L., Arteaga-Vazquez, M., Berger, F., Briginshaw, L.N., Carella, P., Aguilar-Cruz, A., Davies, K.M., Dierschke, T., Dolan, L., Dorantes-Acosta, A.E., Fisher, T.J., Flores-Sandoval, E., Futagami, K., Ishizaki, K., Jibran, R., Kanazawa, T., Kato, H., Kohchi, T., Levins, J., Lin, S.S., Nakagami, H., Nishihama, R., Romani, F., Schornack, S., Tanizawa, Y., Tsuzuki, M., Ueda, T., Watanabe, Y., Yamato, K.T., and Zachgo, S. (2022). The renaissance and enlightenment of Marchantia as a model system. Plant Cell 34, 3512–3542.

Bowman, J.L., Kohchi, T., Yamato, K.T., Jenkins, J., Shu, S., Ishizaki, K., Yamaoka, S., Nishihama, R., Nakamura, Y., Berger, F., Adam, C., Aki, S.S., Althoff, F., Araki, T., Arteaga-Vazquez, M.A., Balasubrmanian, S., Barry, K., Bauer, D., Boehm, C.R., Briginshaw, L., Caballero-Perez, J., Catarino, B., Chen, F., Chiyoda, S., Chovatia, M., Davies, K.M., Delmans, M., Demura, T., Dierschke, T., Dolan, L., Dorantes-Acosta, A.E., Eklund, D.M., Florent, S.N., Flores-Sandoval, E., Fujiyama, A., Fukuzawa, H., Galik, B., Grimanelli, D., Grimwood, J., Grossniklaus, U., Hamada, T., Haseloff, J., Hetherington, A.J., Higo, A., Hirakawa, Y., Hundley, H.N., Ikeda, Y., Inoue, K., Inoue, S.I., Ishida, S., Jia, Q., Kakita, M., Kanazawa, T., Kawai, Y., Kawashima, T., Kennedy, M., Kinose, K., Kinoshita, T., Kohara, Y., Koide, E., Komatsu, K., Kopischke, S., Kubo, M., Kyozuka, J., Lagercrantz, U., Lin, S.S., Lindquist, E., Lipzen, A.M., Lu, C.W., De Luna, E., Martienssen, R.A., Minamino, N., Mizutani, M., Mizutani, M., Mochizuki, N., Monte, I., Mosher, R., Nagasaki, H., Nakagami, H., Naramoto, S., Nishitani, K., Ohtani, M., Okamoto, T., Okumura, M., Phillips, J., Pollak, B., Reinders, A., Rovekamp, M., Sano, R., Sawa, S., Schmid, M.W., Shirakawa, M., Solano, R., Spunde, A., Suetsugu, N., Sugano, S., Sugiyama, A., Sun, R., Suzuki, Y., Takenaka, M., Takezawa, D., Tomogane, H., Tsuzuki, M., Ueda, T., Umeda, M., Ward, J.M., Watanabe, Y., Yazaki, K., Yokoyama, R., Yoshitake, Y., Yotsui, I., Zachgo, S., and Schmutz, J. (2017). Insights into Land Plant Evolution Garnered from the Marchantia polymorpha Genome. Cell 171, 287–304 e215.

Breker, M., Lieberman, K., and Cross, F.R. (2018). Comprehensive Discovery of Cell-Cycle-Essential Pathways in Chlamydomonas reinhardtii. Plant Cell 30, 1178–1198.

Brophy, J.A.N., LaRue, T., and Dinneny, J.R. (2018). Understanding and engineering plant form. Semin Cell Dev Biol 79, 68–77.

Charif, D., and Lobry, J.R. (2007). SeqinR 1.0-2: A Contributed Package to the R Project for Statistical Computing Devoted to Biological Sequences Retrieval and Analysis. In Structural Approaches to Sequence Evolution: Molecules, Networks, Populations, U. Bastolla, M. Porto, H.E. Roman, and M. Vendruscolo, eds (Berlin, Heidelberg: Springer Berlin Heidelberg), pp. 207–232.

Cheng, S., Xian, W., Fu, Y., Marin, B., Keller, J., Wu, T., Sun, W., Li, X., Xu, Y., Zhang, Y., Wittek, S., Reder, T., Gunther, G., Gontcharov, A., Wang, S., Li, L., Liu, X., Wang, J., Yang, H., Xu, X., Delaux, P.M., Melkonian, B., Wong, G.K., and Melkonian, M. (2019). Genomes of Subaerial Zygnematophyceae Provide Insights into Land Plant Evolution. Cell 179, 1057–1067 e1014.

Clute, P., and Pines, J. (1999). Temporal and spatial control of cyclin B1 destruction in metaphase. Nat Cell Biol 1, 82–87.

Cock, J.M., Sterck, L., Rouze, P., Scornet, D., Allen, A.E., Amoutzias, G., Anthouard, V., Artiguenave, F., Aury, J.M., Badger, J.H., Beszteri, B., Billiau, K., Bonnet, E., Bothwell, J.H., Bowler, C., Boyen, C., Brownlee, C., Carrano, C.J., Charrier, B., Cho, G.Y., Coelho, S.M., Collen, J., Corre, E., Da Silva, C., Delage, L., Delaroque, N., Dittami, S.M., Doulbeau, S., Elias, M., Farnham, G., Gachon, C.M., Gschloessl, B., Heesch, S., Jabbari, K., Jubin, C., Kawai, H., Kimura, K., Kloareg, B., Kupper, F.C., Lang, D., Le Bail, A., Leblanc, C., Lerouge, P., Lohr, M., Lopez, P.J., Martens, C., Maumus, F., Michel, G., Miranda-Saavedra, D., Morales, J., Moreau, H., Motomura, T., Nagasato, C., Napoli, C.A., Nelson, D.R., Nyvall-Collen, P., Peters, A.F., Pommier, C., Potin, P., Poulain, J., Quesneville, H., Read, B., Rensing, S.A., Ritter, A., Rousvoal, S., Samanta, M., Samson, G., Schroeder, D.C., Segurens, B., Strittmatter, M., Tonon, T., Tregear, J.W., Valentin, K., von Dassow, P., Yamagishi, T., Van de Peer, Y., and Wincker, P. (2010). The Ectocarpus genome and the independent evolution of multicellularity in brown algae. Nature 465, 617–621.

Consortium, C.e.S. (1998). Genome sequence of the nematode C. elegans: a platform for investigating biology. Science 282, 2012–2018.

Cross, F.R. (2020). Regulation of Multiple Fission and Cell-Cycle-Dependent Gene Expression by CDKA1 and the Rb-E2F Pathway in Chlamydomonas. Curr Biol 30, 1855–1865 e1854.

Cross, F.R., and Umen, J.G. (2015). The Chlamydomonas cell cycle. Plant J 82, 370–392.

d’Erfurth, I., Cromer, L., Jolivet, S., Girard, C., Horlow, C., Sun, Y., To, J.P., Berchowitz, L.E., Copenhaver, G.P., and Mercier, R. (2010). The cyclin-A CYCA1;2/TAM is required for the meiosis I to meiosis II transition and cooperates with OSD1 for the prophase to first meiotic division transition. PLoS Genet 6, e1000989.

Datar, S.A., Jacobs, H.W., de la Cruz, A.F., Lehner, C.F., and Edgar, B.A. (2000). The Drosophila cyclin D-Cdk4 complex promotes cellular growth. EMBO J 19, 4543–4554.

Dewitte, W., Riou-Khamlichi, C., Scofield, S., Healy, J.M., Jacqmard, A., Kilby, N.J., and Murray, J.A. (2003). Altered cell cycle distribution, hyperplasia, and inhibited differentiation in Arabidopsis caused by the D-type cyclin CYCD3. Plant Cell 15, 79–92.

Doerner, P., Jorgensen, J.E., You, R., Steppuhn, J., and Lamb, C. (1996). Control of root growth and development by cyclin expression. Nature 380, 520–523.

Doonan, J., and Hunt, T. (1996). Cell cycle. Why don’t plants get cancer? Nature 380, 481–482.

Feng, G., Burleigh, J.G., Braun, E.L., Mei, W., and Barbazuk, W.B. (2017). Evolution of the 3R-MYB Gene Family in Plants. Genome Biol Evol 9, 1013–1029.

Feng, X., Zheng, J., Irisarri, I., Yu, H., Zheng, B., Ali, Z., de Vries, S., Keller, J., Furst-Jansen, J.M.R., Dadras, A., Zegers, J.M.S., Rieseberg, T.P., Dhabalia Ashok, A., Darienko, T., Bierenbroodspot, M.J., Gramzow, L., Petroll, R., Haas, F.B., Fernandez-Pozo, N., Nousias, O., Li, T., Fitzek, E., Grayburn, W.S., Rittmeier, N., Permann, C., Rumpler, F., Archibald, J.M., Theissen, G., Mower, J.P., Lorenz, M., Buschmann, H., von Schwartzenberg, K., Boston, L., Hayes, R.D., Daum, C., Barry, K., Grigoriev, I.V., Wang, X., Li, F.W., Rensing, S.A., Ben Ari, J., Keren, N., Mosquna, A., Holzinger, A., Delaux, P.M., Zhang, C., Huang, J., Mutwil, M., de Vries, J., and Yin, Y. (2024). Genomes of multicellular algal sisters to land plants illuminate signaling network evolution. Nat Genet 56, 1018–1031.

Finn, R.D., Clements, J., and Eddy, S.R. (2011). HMMER web server: interactive sequence similarity searching. Nucleic Acids Res 39, W29–37.

Geley, S., Kramer, E., Gieffers, C., Gannon, J., Peters, J.M., and Hunt, T. (2001). Anaphase-promoting complex/cyclosome-dependent proteolysis of human cyclin A starts at the beginning of mitosis and is not subject to the spindle assembly checkpoint. J Cell Biol 153, 137–148.

Goffeau, A., Barrell, B.G., Bussey, H., Davis, R.W., Dujon, B., Feldmann, H., Galibert, F., Hoheisel, J.D., Jacq, C., Johnston, M., Louis, E.J., Mewes, H.W., Murakami, Y., Philippsen, P., Tettelin, H., and Oliver, S.G. (1996). Life with 6000 genes. Science 274, 546, 563–547.

Gutierrez, C. (2009). The Arabidopsis cell division cycle. Arabidopsis Book 7, e0120.

Gutierrez, C. (2022). A Journey to the Core of the Plant Cell Cycle. Int J Mol Sci 23.

Hao, Y., Stuart, T., Kowalski, M.H., Choudhary, S., Hoffman, P., Hartman, A., Srivastava, A., Molla, G., Madad, S., Fernandez-Granda, C., and Satija, R. (2024). Dictionary learning for integrative, multimodal and scalable single-cell analysis. Nat Biotechnol 42, 293–304.

Hori, K., Maruyama, F., Fujisawa, T., Togashi, T., Yamamoto, N., Seo, M., Sato, S., Yamada, T., Mori, H., Tajima, N., Moriyama, T., Ikeuchi, M., Watanabe, M., Wada, H., Kobayashi, K., Saito, M., Masuda, T., Sasaki-Sekimoto, Y., Mashiguchi, K., Awai, K., Shimojima, M., Masuda, S., Iwai, M., Nobusawa, T., Narise, T., Kondo, S., Saito, H., Sato, R., Murakawa, M., Ihara, Y., Oshima-Yamada, Y., Ohtaka, K., Satoh, M., Sonobe, K., Ishii, M., Ohtani, R., Kanamori-Sato, M., Honoki, R., Miyazaki, D., Mochizuki, H., Umetsu, J., Higashi, K., Shibata, D., Kamiya, Y., Sato, N., Nakamura, Y., Tabata, S., Ida, S., Kurokawa, K., and Ohta, H. (2014). Klebsormidium flaccidum genome reveals primary factors for plant terrestrial adaptation. Nat Commun 5, 3978.

Imai, K.K., Ohashi, Y., Tsuge, T., Yoshizumi, T., Matsui, M., Oka, A., and Aoyama, T. (2006). The A-type cyclin CYCA2;3 is a key regulator of ploidy levels in Arabidopsis endoreduplication. Plant Cell 18, 382–396.

Inze, D., and De Veylder, L. (2006). Cell cycle regulation in plant development. Annu Rev Genet 40, 77–105.

Ishida, S., Suzuki, H., Iwaki, A., Kawamura, S., Yamaoka, S., Kojima, M., Takebayashi, Y., Yamaguchi, K., Shigenobu, S., Sakakibara, H., Kohchi, T., and Nishihama, R. (2022). Diminished Auxin Signaling Triggers Cellular Reprogramming by Inducing a Regeneration Factor in the Liverwort Marchantia polymorpha. Plant Cell Physiol 63, 384–400.

Ishikawa, M., Murata, T., Sato, Y., Nishiyama, T., Hiwatashi, Y., Imai, A., Kimura, M., Sugimoto, N., Akita, A., Oguri, Y., Friedman, W.E., Hasebe, M., and Kubo, M. (2011). Physcomitrella cyclin-dependent kinase A links cell cycle reactivation to other cellular changes during reprogramming of leaf cells. Plant Cell 23, 2924–2938.

Ito, M., Araki, S., Matsunaga, S., Itoh, T., Nishihama, R., Machida, Y., Doonan, J.H., and Watanabe, A. (2001). G2/M-phase-specific transcription during the plant cell cycle is mediated by c-Myb-like transcription factors. Plant Cell 13, 1891–1905.

Jiao, C., Sorensen, I., Sun, X., Sun, H., Behar, H., Alseekh, S., Philippe, G., Palacio Lopez, K., Sun, L., Reed, R., Jeon, S., Kiyonami, R., Zhang, S., Fernie, A.R., Brumer, H., Domozych, D.S., Fei, Z., and Rose, J.K.C. (2020). The Penium margaritaceum Genome: Hallmarks of the Origins of Land Plants. Cell 181, 1097–1111 e1012.

Joubes, J., Chevalier, C., Dudits, D., Heberle-Bors, E., Inze, D., Umeda, M., and Renaudin, J.P. (2000). CDK-related protein kinases in plants. Plant Mol Biol 43, 607–620.

Katoh, K., Misawa, K., Kuma, K., and Miyata, T. (2002). MAFFT: a novel method for rapid multiple sequence alignment based on fast Fourier transform. Nucleic Acids Res 30, 3059–3066.

Kawamura, S., Romani, F., Yagura, M., Mochizuki, T., Sakamoto, M., Yamaoka, S., Nishihama, R., Nakamura, Y., Yamato, K.T., Bowman, J.L., Kohchi, T., and Tanizawa, Y. (2022). MarpolBase Expression: A Web-Based, Comprehensive Platform for Visualization and Analysis of Transcriptomes in the Liverwort Marchantia polymorpha. Plant Cell Physiol 63, 1745–1755.

Kim, D., Paggi, J.M., Park, C., Bennett, C., and Salzberg, S.L. (2019). Graph-based genome alignment and genotyping with HISAT2 and HISAT-genotype. Nat Biotechnol 37, 907–915.

Kobayashi, K., Suzuki, T., Iwata, E., Magyar, Z., Bogre, L., and Ito, M. (2015). MYB3Rs, plant homologs of Myb oncoproteins, control cell cycle-regulated transcription and form DREAM-like complexes. Transcription 6, 106–111.

Komaki, S., and Sugimoto, K. (2012). Control of the plant cell cycle by developmental and environmental cues. Plant Cell Physiol 53, 953–964.

Koroleva, O.A., Tomlinson, M., Parinyapong, P., Sakvarelidze, L., Leader, D., Shaw, P., and Doonan, J.H. (2004). CycD1, a putative G1 cyclin from Antirrhinum majus, accelerates the cell cycle in cultured tobacco BY-2 cells by enhancing both G1/S entry and progression through S and G2 phases. Plant Cell 16, 2364–2379.

Krylov, D.M., Nasmyth, K., and Koonin, E.V. (2003). Evolution of eukaryotic cell cycle regulation: stepwise addition of regulatory kinases and late advent of the CDKs. Curr Biol 13, 173–177.

Kubota, A., Ishizaki, K., Hosaka, M., and Kohchi, T. (2013). Efficient Agrobacterium-mediated transformation of the liverwort Marchantia polymorpha using regenerating thalli. Biosci Biotechnol Biochem 77, 167–172.

Letunic, I., and Bork, P. (2024). Interactive Tree of Life (iTOL) v6: recent updates to the phylogenetic tree display and annotation tool. Nucleic Acids Res 52, W78–W82.

Li, F.W., Nishiyama, T., Waller, M., Frangedakis, E., Keller, J., Li, Z., Fernandez-Pozo, N., Barker, M.S., Bennett, T., Blazquez, M.A., Cheng, S., Cuming, A.C., de Vries, J., de Vries, S., Delaux, P.M., Diop, I.S., Harrison, C.J., Hauser, D., Hernandez-Garcia, J., Kirbis, A., Meeks, J.C., Monte, I., Mutte, S.K., Neubauer, A., Quandt, D., Robison, T., Shimamura, M., Rensing, S.A., Villarreal, J.C., Weijers, D., Wicke, S., Wong, G.K., Sakakibara, K., and Szovenyi, P. (2020a). Anthoceros genomes illuminate the origin of land plants and the unique biology of hornworts. Nat Plants 6, 259–272.

Li, L., Wang, S., Wang, H., Sahu, S.K., Marin, B., Li, H., Xu, Y., Liang, H., Li, Z., Cheng, S., Reder, T., Cebi, Z., Wittek, S., Petersen, M., Melkonian, B., Du, H., Yang, H., Wang, J., Wong, G.K., Xu, X., Liu, X., Van de Peer, Y., Melkonian, M., and Liu, H. (2020b). The genome of Prasinoderma coloniale unveils the existence of a third phylum within green plants. Nat Ecol Evol 4, 1220–1231.

Liang, Z., Geng, Y., Ji, C., Du, H., Wong, C.E., Zhang, Q., Zhang, Y., Zhang, P., Riaz, A., Chachar, S., Ding, Y., Wen, J., Wu, Y., Wang, M., Zheng, H., Wu, Y., Demko, V., Shen, L., Han, X., Zhang, P., Gu, X., and Yu, H. (2020). Mesostigma viride Genome and Transcriptome Provide Insights into the Origin and Evolution of Streptophyta. Adv Sci (Weinh) 7, 1901850.

Lifschitz, S., Haeusler, E.H., Catanho, M., Miranda, A.B., Armas, E.M., Heine, A., Moreira, S., and Tristao, C. (2022). Bio-Strings: A Relational Database Data-Type for Dealing with Large Biosequences. BioTech (Basel) 11.

Linde, A.M., Sawangproh, W., Cronberg, N., Szovenyi, P., and Lagercrantz, U. (2020). Evolutionary History of the Marchantia polymorpha Complex. Front Plant Sci 11, 829.

Littlejohn, G.R., Gouveia, J.D., Edner, C., Smirnoff, N., and Love, J. (2010). Perfluorodecalin enhances in vivo confocal microscopy resolution of Arabidopsis thaliana mesophyll. New Phytol 186, 1018–1025.

Liu, J., and Kipreos, E.T. (2000). Evolution of cyclin-dependent kinases (CDKs) and CDK-activating kinases (CAKs): differential conservation of CAKs in yeast and metazoa. Mol Biol Evol 17, 1061–1074.

Liu, W., Cai, G., Zhai, N., Wang, H., Tang, T., Zhang, Y., Zhang, Z., Sun, L., Zhang, Y., Beeckman, T., and Xu, L. (2023). Genome and transcriptome of Selaginella kraussiana reveal evolution of root apical meristems in vascular plants. Curr Biol 33, 4085–4097 e4085.

Ma, Z., Wu, Y., Jin, J., Yan, J., Kuang, S., Zhou, M., Zhang, Y., and Guo, A.Y. (2013). Phylogenetic analysis reveals the evolution and diversification of cyclins in eukaryotes. Mol Phylogenet Evol 66, 1002–1010.

Marchant, D.B., Chen, G., Cai, S., Chen, F., Schafran, P., Jenkins, J., Shu, S., Plott, C., Webber, J., Lovell, J.T., He, G., Sandor, L., Williams, M., Rajasekar, S., Healey, A., Barry, K., Zhang, Y., Sessa, E., Dhakal, R.R., Wolf, P.G., Harkess, A., Li, F.W., Rossner, C., Becker, A., Gramzow, L., Xue, D., Wu, Y., Tong, T., Wang, Y., Dai, F., Hua, S., Wang, H., Xu, S., Xu, F., Duan, H., Theissen, G., McKain, M.R., Li, Z., McKibben, M.T.W., Barker, M.S., Schmitz, R.J., Stevenson, D.W., Zumajo-Cardona, C., Ambrose, B.A., Leebens-Mack, J.H., Grimwood, J., Schmutz, J., Soltis, P.S., Soltis, D.E., and Chen, Z.H. (2022). Dynamic genome evolution in a model fern. Nat Plants 8, 1038–1051.

Martinez-Alonso, D., and Malumbres, M. (2020). Mammalian cell cycle cyclins. Semin Cell Dev Biol 107, 28–35.

Matsuzaki, M., Misumi, O., Shin, I.T., Maruyama, S., Takahara, M., Miyagishima, S.Y., Mori, T., Nishida, K., Yagisawa, F., Nishida, K., Yoshida, Y., Nishimura, Y., Nakao, S., Kobayashi, T., Momoyama, Y., Higashiyama, T., Minoda, A., Sano, M., Nomoto, H., Oishi, K., Hayashi, H., Ohta, F., Nishizaka, S., Haga, S., Miura, S., Morishita, T., Kabeya, Y., Terasawa, K., Suzuki, Y., Ishii, Y., Asakawa, S., Takano, H., Ohta, N., Kuroiwa, H., Tanaka, K., Shimizu, N., Sugano, S., Sato, N., Nozaki, H., Ogasawara, N., Kohara, Y., and Kuroiwa, T. (2004). Genome sequence of the ultrasmall unicellular red alga Cyanidioschyzon merolae 10D. Nature 428, 653–657.

Menges, M., de Jager, S.M., Gruissem, W., and Murray, J.A. (2005). Global analysis of the core cell cycle regulators of Arabidopsis identifies novel genes, reveals multiple and highly specific profiles of expression and provides a coherent model for plant cell cycle control. Plant J 41, 546–566.

Menges, M., Samland, A.K., Planchais, S., and Murray, J.A. (2006). The D-type cyclin CYCD3;1 is limiting for the G1-to-S-phase transition in Arabidopsis. Plant Cell 18, 893–906.

Merchant, S.S., Prochnik, S.E., Vallon, O., Harris, E.H., Karpowicz, S.J., Witman, G.B., Terry, A., Salamov, A., Fritz-Laylin, L.K., Marechal-Drouard, L., Marshall, W.F., Qu, L.H., Nelson, D.R., Sanderfoot, A.A., Spalding, M.H., Kapitonov, V.V., Ren, Q., Ferris, P., Lindquist, E., Shapiro, H., Lucas, S.M., Grimwood, J., Schmutz, J., Cardol, P., Cerutti, H., Chanfreau, G., Chen, C.L., Cognat, V., Croft, M.T., Dent, R., Dutcher, S., Fernandez, E., Fukuzawa, H., Gonzalez-Ballester, D., Gonzalez-Halphen, D., Hallmann, A., Hanikenne, M., Hippler, M., Inwood, W., Jabbari, K., Kalanon, M., Kuras, R., Lefebvre, P.A., Lemaire, S.D., Lobanov, A.V., Lohr, M., Manuell, A., Meier, I., Mets, L., Mittag, M., Mittelmeier, T., Moroney, J.V., Moseley, J., Napoli, C., Nedelcu, A.M., Niyogi, K., Novoselov, S.V., Paulsen, I.T., Pazour, G., Purton, S., Ral, J.P., Riano-Pachon, D.M., Riekhof, W., Rymarquis, L., Schroda, M., Stern, D., Umen, J., Willows, R., Wilson, N., Zimmer, S.L., Allmer, J., Balk, J., Bisova, K., Chen, C.J., Elias, M., Gendler, K., Hauser, C., Lamb, M.R., Ledford, H., Long, J.C., Minagawa, J., Page, M.D., Pan, J., Pootakham, W., Roje, S., Rose, A., Stahlberg, E., Terauchi, A.M., Yang, P., Ball, S., Bowler, C., Dieckmann, C.L., Gladyshev, V.N., Green, P., Jorgensen, R., Mayfield, S., Mueller-Roeber, B., Rajamani, S., Sayre, R.T., Brokstein, P., Dubchak, I., Goodstein, D., Hornick, L., Huang, Y.W., Jhaveri, J., Luo, Y., Martinez, D., Ngau, W.C., Otillar, B., Poliakov, A., Porter, A., Szajkowski, L., Werner, G., Zhou, K., Grigoriev, I.V., Rokhsar, D.S., and Grossman, A.R. (2007). The Chlamydomonas genome reveals the evolution of key animal and plant functions. Science 318, 245–250.

Minh, B.Q., Schmidt, H.A., Chernomor, O., Schrempf, D., Woodhams, M.D., von Haeseler, A., and Lanfear, R. (2020). IQ-TREE 2: New Models and Efficient Methods for Phylogenetic Inference in the Genomic Era. Mol Biol Evol 37, 1530–1534.

Mironov, V.V., De Veylder, L., Van Montagu, M., and Inze, D. (1999). Cyclin-dependent kinases and cell division in plants-the nexus. Plant Cell 11, 509–522.

Montgomery, S.A., Tanizawa, Y., Galik, B., Wang, N., Ito, T., Mochizuki, T., Akimcheva, S., Bowman, J.L., Cognat, V., Marechal-Drouard, L., Ekker, H., Hong, S.F., Kohchi, T., Lin, S.S., Liu, L.D., Nakamura, Y., Valeeva, L.R., Shakirov, E.V., Shippen, D.E., Wei, W.L., Yagura, M., Yamaoka, S., Yamato, K.T., Liu, C., and Berger, F. (2020). Chromatin Organization in Early Land Plants Reveals an Ancestral Association between H3K27me3, Transposons, and Constitutive Heterochromatin. Curr Biol 30, 573–588 e577.

Nagai, I. (1919). Induced Adventitious Growth in the Gemnnae of Marchanti. The Botanical Magazine 33, 99–109.

Naito, Y., Hino, K., Bono, H., and Ui-Tei, K. (2015). CRISPRdirect: software for designing CRISPR/Cas guide RNA with reduced off-target sites. Bioinformatics 31, 1120–1123.

Nishihama, R., Ishida, S., Urawa, H., Kamei, Y., and Kohchi, T. (2016). Conditional Gene Expression/Deletion Systems for Marchantia polymorpha Using its Own Heat-Shock Promoter and Cre/loxP-Mediated Site-Specific Recombination. Plant Cell Physiol 57, 271–280.

Nishihama, R., Ishizaki, K., Hosaka, M., Matsuda, Y., Kubota, A., and Kohchi, T. (2015). Phytochrome-mediated regulation of cell division and growth during regeneration and sporeling development in the liverwort Marchantia polymorpha. J Plant Res 128, 407–421.

Nishiyama, T., Sakayama, H., de Vries, J., Buschmann, H., Saint-Marcoux, D., Ullrich, K.K., Haas, F.B., Vanderstraeten, L., Becker, D., Lang, D., Vosolsobe, S., Rombauts, S., Wilhelmsson, P.K.I., Janitza, P., Kern, R., Heyl, A., Rumpler, F., Villalobos, L., Clay, J.M., Skokan, R., Toyoda, A., Suzuki, Y., Kagoshima, H., Schijlen, E., Tajeshwar, N., Catarino, B., Hetherington, A.J., Saltykova, A., Bonnot, C., Breuninger, H., Symeonidi, A., Radhakrishnan, G.V., Van Nieuwerburgh, F., Deforce, D., Chang, C., Karol, K.G., Hedrich, R., Ulvskov, P., Glockner, G., Delwiche, C.F., Petrasek, J., Van de Peer, Y., Friml, J., Beilby, M., Dolan, L., Kohara, Y., Sugano, S., Fujiyama, A., Delaux, P.M., Quint, M., Theissen, G., Hagemann, M., Harholt, J., Dunand, C., Zachgo, S., Langdale, J., Maumus, F., Van Der Straeten, D., Gould, S.B., and Rensing, S.A. (2018). The Chara Genome: Secondary Complexity and Implications for Plant Terrestrialization. Cell 174, 448–464 e424.

Nowack, M.K., Harashima, H., Dissmeyer, N., Zhao, X., Bouyer, D., Weimer, A.K., De Winter, F., Yang, F., and Schnittger, A. (2012). Genetic framework of cyclin-dependent kinase function in Arabidopsis. Dev Cell 22, 1030–1040.

Palenik, B., Grimwood, J., Aerts, A., Rouze, P., Salamov, A., Putnam, N., Dupont, C., Jorgensen, R., Derelle, E., Rombauts, S., Zhou, K., Otillar, R., Merchant, S.S., Podell, S., Gaasterland, T., Napoli, C., Gendler, K., Manuell, A., Tai, V., Vallon, O., Piganeau, G., Jancek, S., Heijde, M., Jabbari, K., Bowler, C., Lohr, M., Robbens, S., Werner, G., Dubchak, I., Pazour, G.J., Ren, Q., Paulsen, I., Delwiche, C., Schmutz, J., Rokhsar, D., Van de Peer, Y., Moreau, H., and Grigoriev, I.V. (2007). The tiny eukaryote Ostreococcus provides genomic insights into the paradox of plankton speciation. Proc Natl Acad Sci U S A 104, 7705–7710.

Patron, N.J., Orzaez, D., Marillonnet, S., Warzecha, H., Matthewman, C., Youles, M., Raitskin, O., Leveau, A., Farre, G., Rogers, C., Smith, A., Hibberd, J., Webb, A.A., Locke, J., Schornack, S., Ajioka, J., Baulcombe, D.C., Zipfel, C., Kamoun, S., Jones, J.D., Kuhn, H., Robatzek, S., Van Esse, H.P., Sanders, D., Oldroyd, G., Martin, C., Field, R., O’Connor, S., Fox, S., Wulff, B., Miller, B., Breakspear, A., Radhakrishnan, G., Delaux, P.M., Loque, D., Granell, A., Tissier, A., Shih, P., Brutnell, T.P., Quick, W.P., Rischer, H., Fraser, P.D., Aharoni, A., Raines, C., South, P.F., Ane, J.M., Hamberger, B.R., Langdale, J., Stougaard, J., Bouwmeester, H., Udvardi, M., Murray, J.A., Ntoukakis, V., Schafer, P., Denby, K., Edwards, K.J., Osbourn, A., and Haseloff, J. (2015). Standards for plant synthetic biology: a common syntax for exchange of DNA parts. New Phytol 208, 13–19.

Pecani, K., Lieberman, K., Tajima-Shirasaki, N., Onishi, M., and Cross, F.R. (2022). Control of division in Chlamydomonas by cyclin B/CDKB1 and the anaphase-promoting complex. PLoS Genet 18, e1009997.

Peramuna, A., Lopez, C.Q., Rios, F.J.A., Bae, H., Fangel, J.U., Batth, R., Harholt, J., and Simonsen, H.T. (2023). Overexpression of Physcomitrium patens cell cycle regulators leads to larger gametophytes. Sci Rep 13, 4301.

Procter, J.B., Carstairs, G.M., Soares, B., Mourao, K., Ofoegbu, T.C., Barton, D., Lui, L., Menard, A., Sherstnev, N., Roldan-Martinez, D., Duce, S., Martin, D.M.A., and Barton, G.J. (2021). Alignment of Biological Sequences with Jalview. Methods Mol Biol 2231, 203–224.

Quevillon, E., Silventoinen, V., Pillai, S., Harte, N., Mulder, N., Apweiler, R., and Lopez, R. (2005). InterProScan: protein domains identifier. Nucleic Acids Res 33, W116–120.

Radhakrishnan, G.V., Keller, J., Rich, M.K., Vernie, T., Mbadinga Mbadinga, D.L., Vigneron, N., Cottret, L., Clemente, H.S., Libourel, C., Cheema, J., Linde, A.M., Eklund, D.M., Cheng, S., Wong, G.K.S., Lagercrantz, U., Li, F.W., Oldroyd, G.E.D., and Delaux, P.M. (2020). An ancestral signalling pathway is conserved in intracellular symbioses-forming plant lineages. Nat Plants 6, 280–289.

Rauber, R., Cabreira, C., de Freitas, L.B., Turchetto-Zolet, A.C., and Margis-Pinheiro, M. (2016). The evolutionary history of the E2F and DEL genes in Viridiplantae. Mol Phylogenet Evol 99, 225–234.

Romani, F., Sauret-Gueto, S., Rebmann, M., Annese, D., Bonter, I., Tomaselli, M., Dierschke, T., Delmans, M., Frangedakis, E., Silvestri, L., Rever, J., Bowman, J.L., Romani, I., and Haseloff, J. (2024). The landscape of transcription factor promoter activity during vegetative development in Marchantia. Plant Cell 36, 2140–2159.

Sakamoto, Y., Ishimoto, A., Sakai, Y., Sato, M., Nishihama, R., Abe, K., Sano, Y., Furuichi, T., Tsuji, H., Kohchi, T., and Matsunaga, S. (2022). Improved clearing method contributes to deep imaging of plant organs. Commun Biol 5, 12.

Sauret-Gueto, S., Frangedakis, E., Silvestri, L., Rebmann, M., Tomaselli, M., Markel, K., Delmans, M., West, A., Patron, N.J., and Haseloff, J. (2020). Systematic Tools for Reprogramming Plant Gene Expression in a Simple Model, Marchantia polymorpha. ACS Synth Biol 9, 864–882.

Schnittger, A., and De Veylder, L. (2018). The Dual Face of Cyclin B1. Trends Plant Sci 23, 475–478.

Schnittger, A., Schobinger, U., Stierhof, Y.D., and Hulskamp, M. (2002a). Ectopic B-type cyclin expression induces mitotic cycles in endoreduplicating Arabidopsis trichomes. Curr Biol 12, 415–420.

Schnittger, A., Schobinger, U., Bouyer, D., Weinl, C., Stierhof, Y.D., and Hulskamp, M. (2002b). Ectopic D-type cyclin expression induces not only DNA replication but also cell division in Arabidopsis trichomes. Proc Natl Acad Sci U S A 99, 6410–6415.

Schween, G., Gorr, G., Hohe, A., and Reski, R. (2008). Unique Tissue-Specific Cell Cycle in Physcomitrella. Plant Biology 5, 50–58.

Siligato, R., Wang, X., Yadav, S.R., Lehesranta, S., Ma, G., Ursache, R., Sevilem, I., Zhang, J., Gorte, M., Prasad, K., Wrzaczek, M., Heidstra, R., Murphy, A., Scheres, B., and Mahonen, A.P. (2016). MultiSite Gateway-Compatible Cell Type-Specific Gene-Inducible System for Plants. Plant Physiol 170, 627–641.

Sozzani, R., Maggio, C., Varotto, S., Canova, S., Bergounioux, C., Albani, D., and Cella, R. (2006). Interplay between Arabidopsis activating factors E2Fb and E2Fa in cell cycle progression and development. Plant Physiol 140, 1355–1366.

Sugano, S.S., Nishihama, R., Shirakawa, M., Takagi, J., Matsuda, Y., Ishida, S., Shimada, T., Hara-Nishimura, I., Osakabe, K., and Kohchi, T. (2018). Efficient CRISPR/Cas9-based genome editing and its application to conditional genetic analysis in Marchantia polymorpha. PLoS One 13, e0205117.

Takahashi, I., Kojima, S., Sakaguchi, N., Umeda-Hara, C., and Umeda, M. (2010). Two Arabidopsis cyclin A3s possess G1 cyclin-like features. Plant Cell Rep 29, 307–315.

Toyoshima, F., Moriguchi, T., Wada, A., Fukuda, M., and Nishida, E. (1998). Nuclear export of cyclin B1 and its possible role in the DNA damage-induced G2 checkpoint. EMBO J 17, 2728–2735.

Tse, S.W., Annese, D., Romani, F., Guzman-Chavez, F., Bonter, I., Forestier, E., Frangedakis, E., and Haseloff, J. (2024). Optimising promoters and subcellular localisation for constitutive transgene expression in Marchantia polymorpha. Plant Cell Physiol.

Tulin, F., and Cross, F.R. (2014). A microbial avenue to cell cycle control in the plant superkingdom. Plant Cell 26, 4019–4038.

Tulin, F., and Cross, F.R. (2015). Cyclin-Dependent Kinase Regulation of Diurnal Transcription in Chlamydomonas. Plant Cell 27, 2727–2742.

Wang, L., Wan, M.C., Liao, R.Y., Xu, J., Xu, Z.G., Xue, H.C., Mai, Y.X., and Wang, J.W. (2023). The maturation and aging trajectory of Marchantia polymorpha at single-cell resolution. Dev Cell.

Wang, W., Haberer, G., Gundlach, H., Glasser, C., Nussbaumer, T., Luo, M.C., Lomsadze, A., Borodovsky, M., Kerstetter, R.A., Shanklin, J., Byrant, D.W., Mockler, T.C., Appenroth, K.J., Grimwood, J., Jenkins, J., Chow, J., Choi, C., Adam, C., Cao, X.H., Fuchs, J., Schubert, I., Rokhsar, D., Schmutz, J., Michael, T.P., Mayer, K.F., and Messing, J. (2014). The Spirodela polyrhiza genome reveals insights into its neotenous reduction fast growth and aquatic lifestyle. Nat Commun 5, 3311.

Willems, A., Heyman, J., Eekhout, T., Achon, I., Pedroza-Garcia, J.A., Zhu, T., Li, L., Vercauteren, I., Van den Daele, H., van de Cotte, B., De Smet, I., and De Veylder, L. (2020). The Cyclin CYCA3;4 Is a Postprophase Target of the APC/C(CCS52A2) E3-Ligase Controlling Formative Cell Divisions in Arabidopsis. Plant Cell 32, 2979–2996.

Zhang, T.Q., Chen, Y., and Wang, J.W. (2021). A single-cell analysis of the Arabidopsis vegetative shoot apex. Dev Cell 56, 1056–1074 e1058.

