## Supplemental Figures for "The minimal cell-cycle control system in *Marchantia* as a framework for understanding plant cell proliferation"

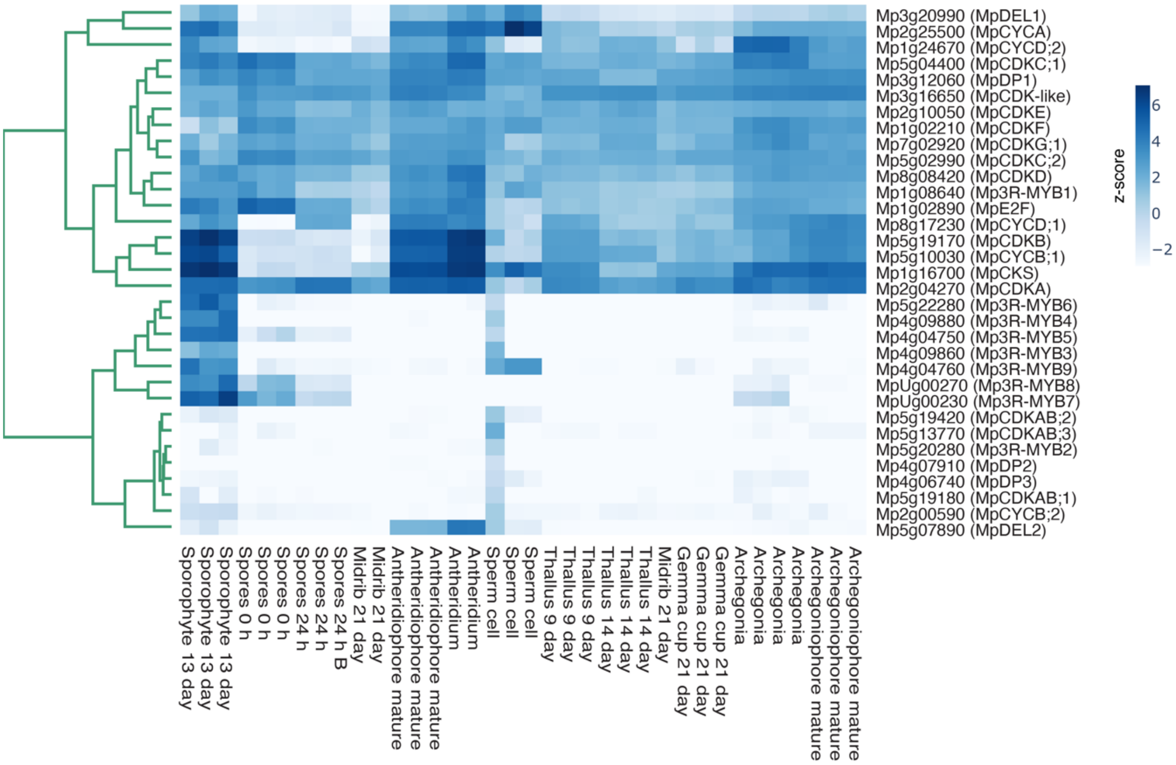


***Suppl. Figure 1. Expression analysis of core cell cycle genes across Marchantia development.*** *RNA-seq analysis of representative cell cycle genes during the time-course of regeneration. Genes are grouped per families and distinguished by colour (see legend). Individual points represent biological replicates and lines averages. b) Heatmap of cell cycle genes across Marchantia tissues during its life cycle as indicated in the x-axis. Colours represent Z-scores of normalized TPM. TPM, transcripts per million.*

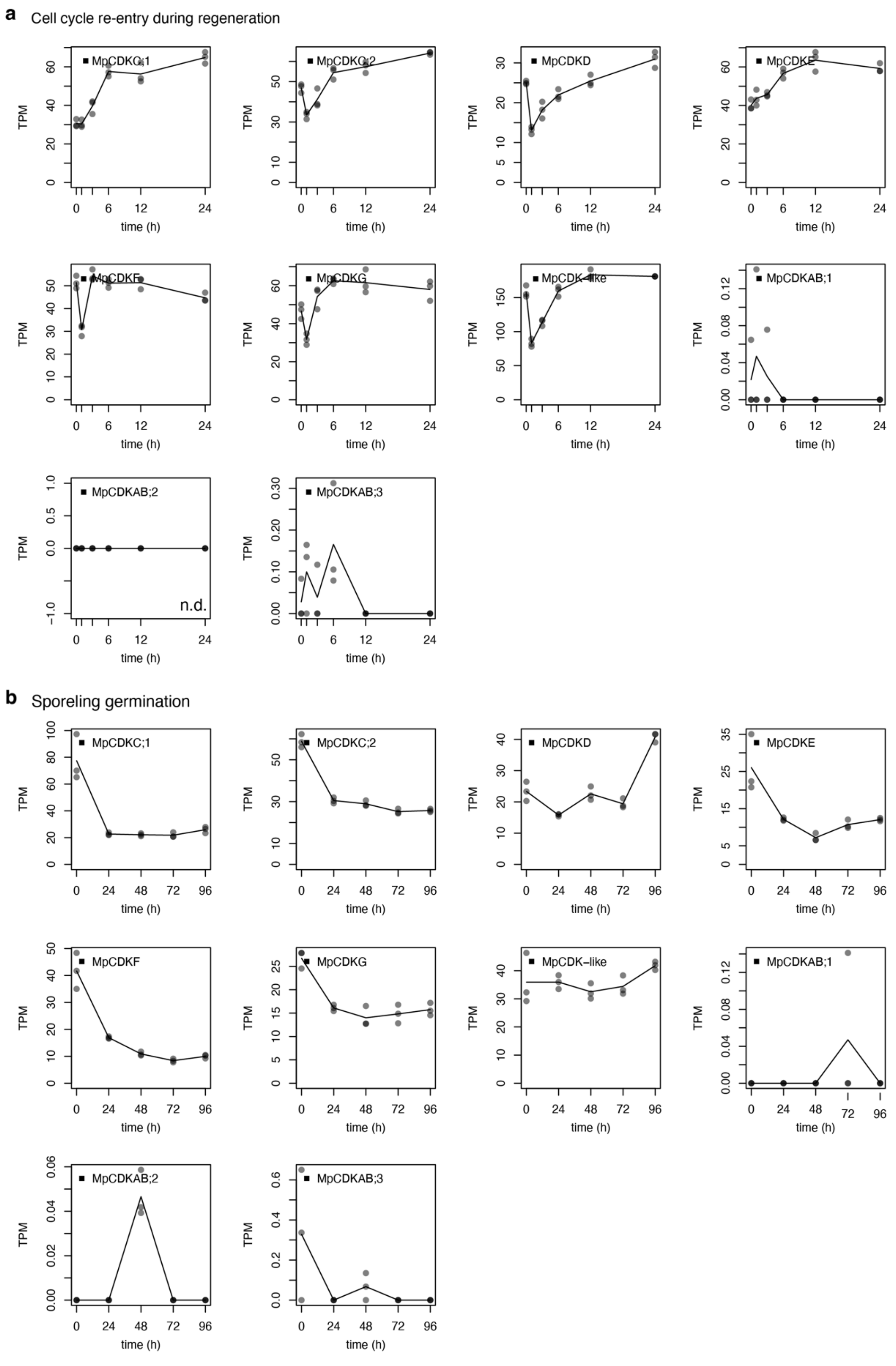


***Suppl. Figure 2.*** **Expression analysis of other CDKs in regeneration and sporeling germination in Marchantia.** *RNA-seq of other CDKs in regeneration and sporeling germination. Individual points represent biological replicates and lines averages. TPM, transcripts per million.*

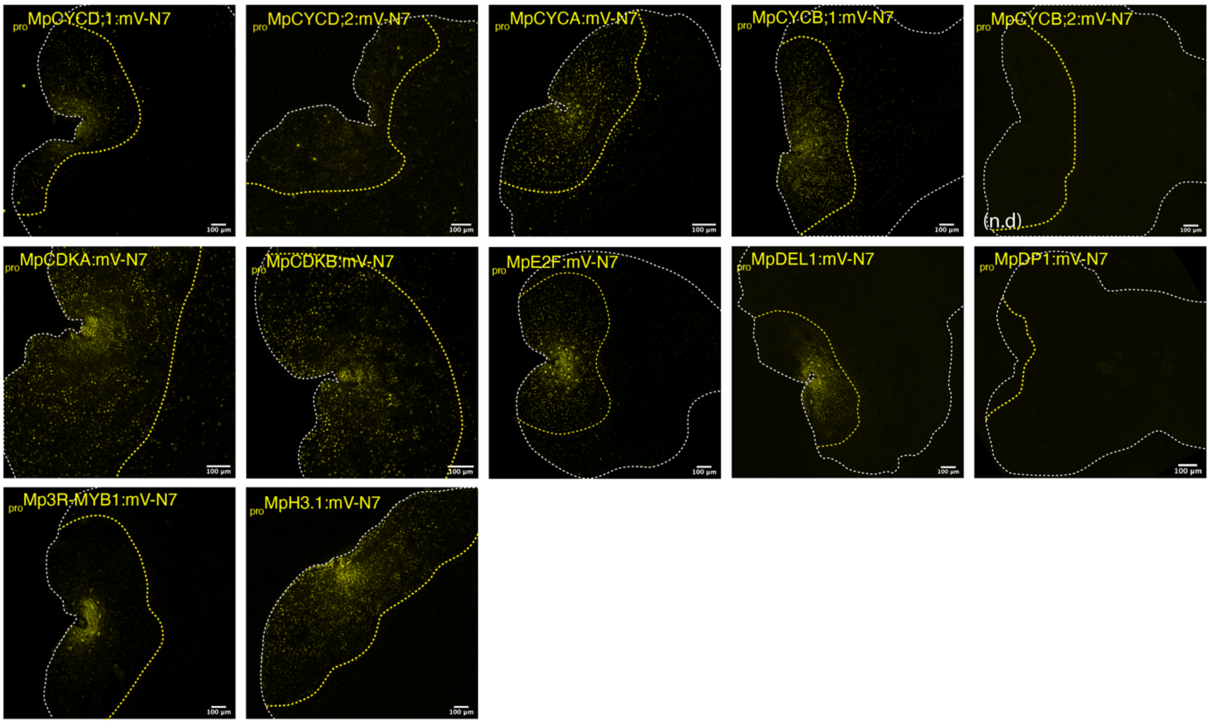


**Suppl. Figure 3. *Transcriptional reporters of selected cell cycle genes.*** *Confocal images of* transcriptional fluorescent reporters (mVenus, yellow) in 5-days old gemmalings. Reporter signal in yellow. *The white dashed lines delimitate the plant outlines. The yellow dashed lines mark the boundary of the mature epidermis.* Scale bar is shown in each image.

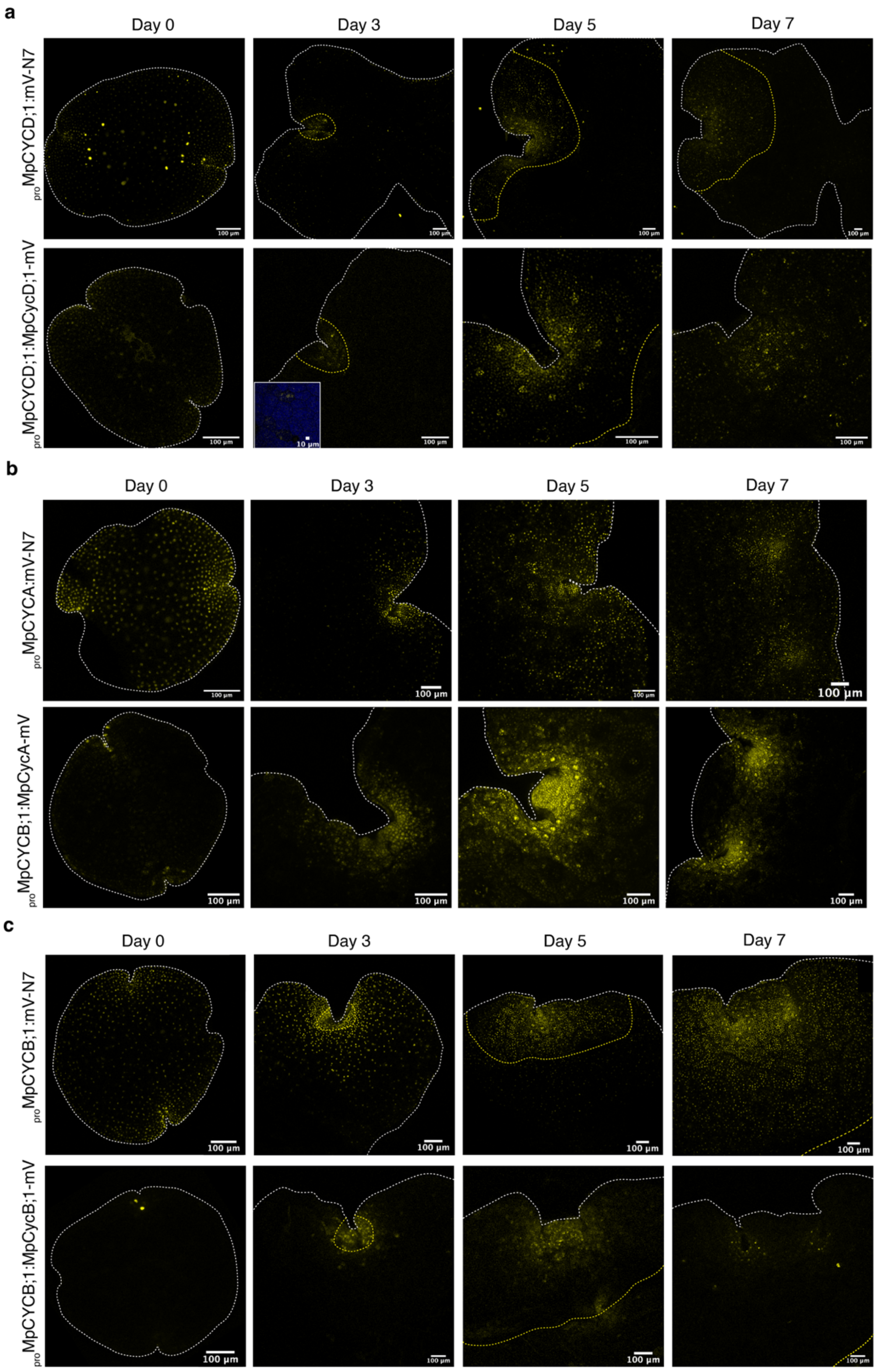

a
Day 0
Day 3
Day 5
Day 7
proMpCYCD;1:mV-N7
proMpCYCD;1:MpCycD;1-mV
100 µm
10 µm
b
Day 0
Day 3
Day 5
Day 7
proMpCYCA:mV-N7
proMpCYCB;1:MpCycA-mV
100 µm
c
Day 0
Day 3
Day 5
Day 7
proMpCYCB;1:mV-N7
proMpCYCB;1:MpCycB;1-mV
100 µm

***Suppl. Figure 4. Time-course of transcriptional reporters of cyclins and CDKs.*** *Time-courses of MpCYCD;1 (a), MpCYCA (b), and MpCYCB;1 (c) transcriptional and translational reporters (mVenus, yellow) in Marchantia gemmallings (0, 3, 5, and 7 days). The white dashed lines delimit the outlines of the plants. The yellow dashed lines mark the boundary of the mature epidermis. Scale bar is shown in each image. The square in MpCYCD;1 highlight a rhizoid precursor cell (chlorophyll autofluorescence in blue).*

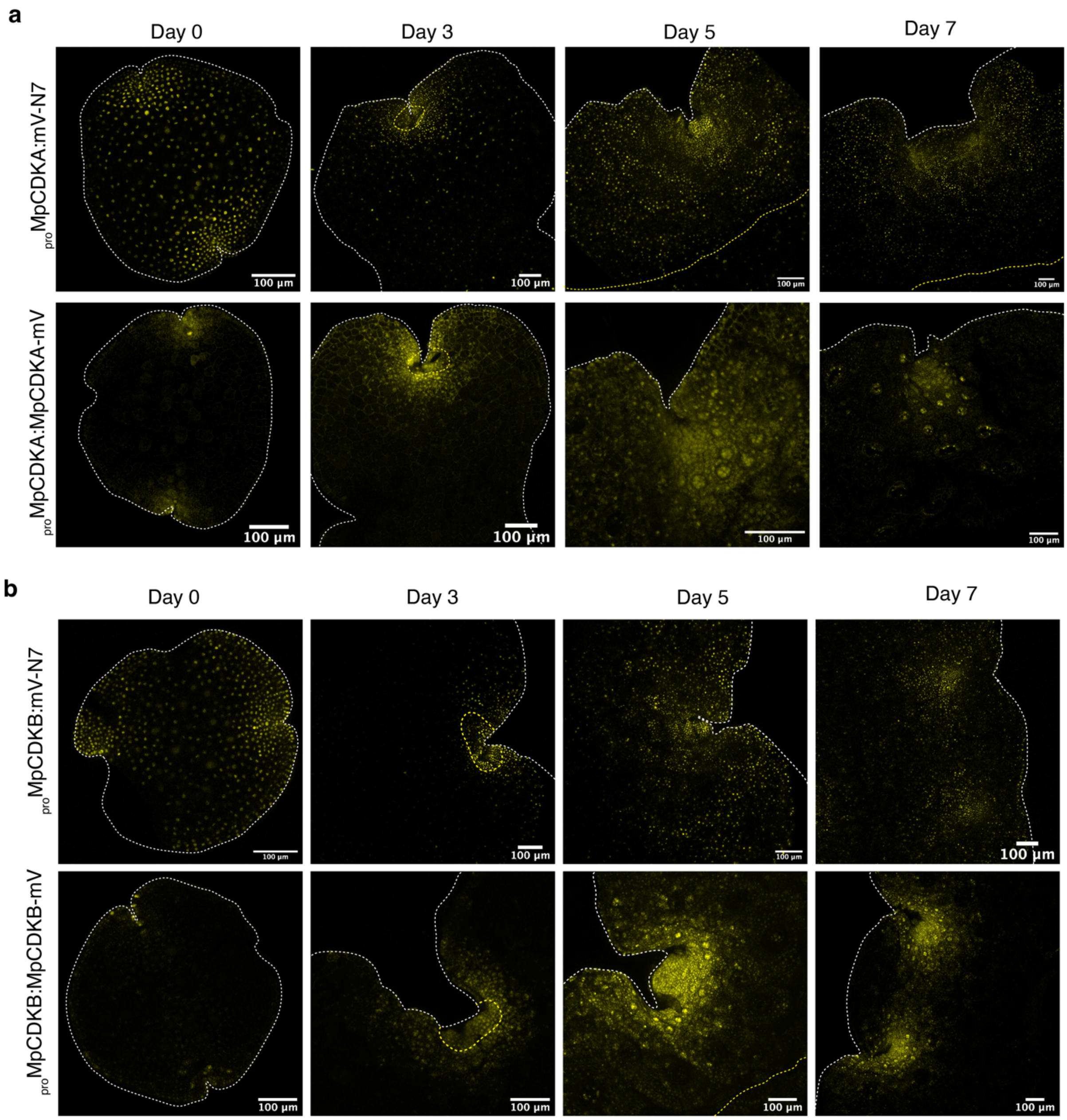


***Suppl. Figure 5. Time-course of translational reporters of cyclins and CDKs.***
*Time-courses of MpCDKA (a), and MpCYCB;1 (b) transcriptional and translational reporters (yellow) in Marchantia gemmallings (0, 3, 5, and 7 days). The white dashed line delimitates the outline of the plant. The yellow dashed line represents the boundary between the mature epidermis and the supportive tissue of the gemmae. Scale bar is shown in each image.*

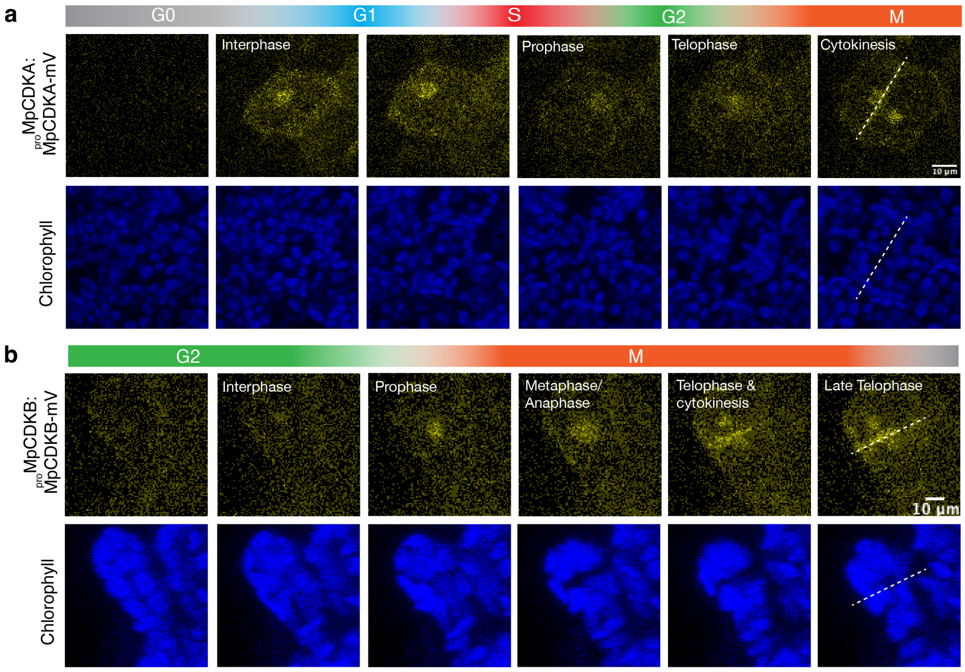


***Suppl. Figure 6. Time-lapse of translational reporters of cyclins and CDKs on individual cells.*** *a-b) Time-lapse of MpCDKA (a) and MpCDKB (b) translational reporters of an individual cell dividing during regeneration (yellow). Chlorophyll (blue) is shown to visualize the cell division phases using chloroplast movement. The full videos of the time-lapses are available at Suppl. Video 4-5.*

a

Mp*1g24670*/Mp*CYCD;2*

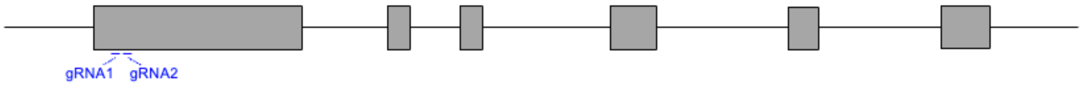


gRNA1

```
WT              CAAGCTTCTGTCCAAGTCCAAGCCGT--------------------CGGGCGGCTCACGA
Mpcycd;2-1ge    CAAGCTTCTGTCCAAGTCCAAGCCGTCATCTCACGGCGGACTCAAGCGGGCGGCTCACGA
```

gRNA2

```
WT              ACCATGCACGCCTGCCCGCTTGCGCTC-CAGAGGCCGTGGA
Mpcycd;2-2ge    ACCATGCACGCCTGCCCGCTTGCGCTCTCAGAGGCCGTGGA
```

b

```
WT
MCVRYFSRRRCQSSKKTFRKKLRPSPECHKLLSKSKPSGGSRDHARLPACAPEAVDLFCPESLPMLDPQSCQVSSDA
AAAAVEPSDLPTDELRLVAPMTTPSSSSSSDPGAAFNAKSPPGCNVKDGVWHDIDGMPQMLPSMGPRAADPRHGLAD
GGGWHNFCPDPGSQPGLGAVKRRGWHLETAASFISSGNSRQSNYLPCERWPVRARAPLSLPSTRTACHDRRQPTKPS
SSFRINGAMRLPDDHDFSCPSKFGEAMGILLEKETASCFQLSKPYIDGLTENSAQKLEHNELRDERRAIVNWIFEVT
RLLQLSSLSAAHAVSFLDRYLSQRFFPDAEFGANVDIRLAATACIWIAAKLEDSGAPDLSSMRELFSLDYDEEKIRK
MEVSVLKCLGWRMLAVTALDFLANMVAQLPAFGLPTHLLRPVGHRSEEIMQGLLSEVEFLAYQPSVVSMSVMQCLLD
ECVPVHTASIMAAFHLALAFDVESEKVCHRTLMDLVVDPDLVCSLPSPDRVDCASSREEPSGCSQWTDAIGHRYLLS
TSTSSSLRSSEILKC*
Mpcycd;2-1ge
MCVRYFSRRRCQSSKKTFRKKLRPSPECHKLLSKSKPSSHGGLKRAAHETMHACPLALQRPWTCFAPSRSPC*
Mpcycd;2-2ge
MCVRYFSRRRCQSSKKTFRKKLRPSPECHKLLSKSKPSGGSRDHARLPACALRGRGLVLPRVAPHVRSTELSGLFRC
CCCSSRAV*
```

***Suppl. Figure 7. Genotyping of Mpcycd;2$^{ge}$.*** *a) Structure of Mp1g24670/MpCYCD;2 locus with the position of designed guide RNA (gRNA). Exons are shown as boxes. Genotyping of genome editing alleles (bottom). gRNA sequence is in blue. Deleted or inserted bases are indicated in red. b) Wild-type (WT) and mutant protein sequences* *deduced from the genomic DNA sequences are indicated. Sequences different from WT are in red. Asterisks indicate translational termination.*

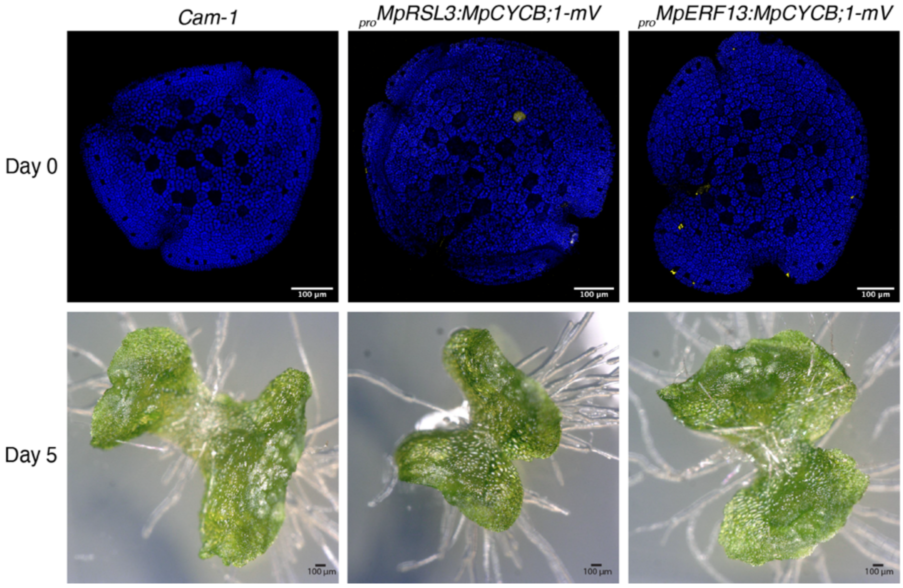


***Suppl. Figure 8. Cell-type specific overexpression of MpCYCB;1 using cell-types specific promoters.*** *Plants expressing $_{pro}$MpRSL3:MpCYCB;1-mVenus (rhizoid precursors specific) and $_{pro}$MpERF13:MpCYCD;1-mVenus (oil bodies specific). Wild-type Cam-1 is used as a control. Confocal full-stack images of representative individual gemma are shown with mVenus (yellow) and chlorophyll (blue) channels merged. Representative picture of 5-days old gemmalings. Scale bar lengths are indicated in the pictures.*

**Supplemental Videos:**

***Suppl. Video 1.*** *$_{pro}$MpCYCD;1:MpCYCD;1-mVenus time-lapse during regeneration. mVenus in yellow and chlorophyll autofluorescence in blue. Scale bar is indicated in the video.*

***Suppl. Video 2.*** *$_{pro}$MpCYCA:MpCYCA-mVenus time-lapse during regeneration. mVenus in yellow and chlorophyll autofluorescence in blue. Scale bar is indicated in the video.*

***Suppl. Video 3.*** *$_{pro}$MpCYCB;1:MpCYCB;1-mVenus time-lapse during regeneration. mVenus in yellow and chlorophyll autofluorescence in blue. Scale bar is indicated in the video.*

***Suppl. Video 4.*** *$_{pro}$MpCDKA:MpCDKA-mVenus time-lapse during regeneration. mVenus in yellow and chlorophyll autofluorescence in blue. Scale bar is indicated in the video.*

***Suppl. Video 5.*** *$_{pro}$MpCDKB:MpCDKB-mVenus time-lapse during regeneration. mVenus in yellow and chlorophyll autofluorescence in blue. Scale bar is indicated in the video.*

***Suppl. Video 6.*** *$_{pro}$35S:MpCYCB1-mVenus time-lapse during gemmallings development. mVenus in yellow. Scale bar is indicated in the video.*

***Suppl. Video 7.*** *$_{pro}$MpCYCB;1:MpCYCB1-mVenus time-lapse during gemmallings development. mVenus in yellow. Scale bar is indicated in the video.*

***Supplemental Tables:***

***Supplemental Table 1.*** *Complete list of scRNA-seq cluster-specific markers and gene annotations.*

**Supplemental Table 2.** Vectors, cloning, primers, and details of gene IDs used.
