## Supplemental File 1 for "The minimal cell-cycle control system in *Marchantia* as a framework for understanding plant cell proliferation"

***Suppl. File 1. Supporting information for phylogenetic analysis of cell-cycle proteins.*** *a) Phylogeny of Cyclins. b) Phylogeny of the CDK and PSTAIRE motifs. c) Phylogeny of CKS proteins. d) Phylogeny of E2F/DP/DEL proteins. e) Phylogeny of RBR. f) Phylogeny of KRM and SMR proteins. g) Phylogeny of 3R-MYBs and analysis of sequence repeats.*

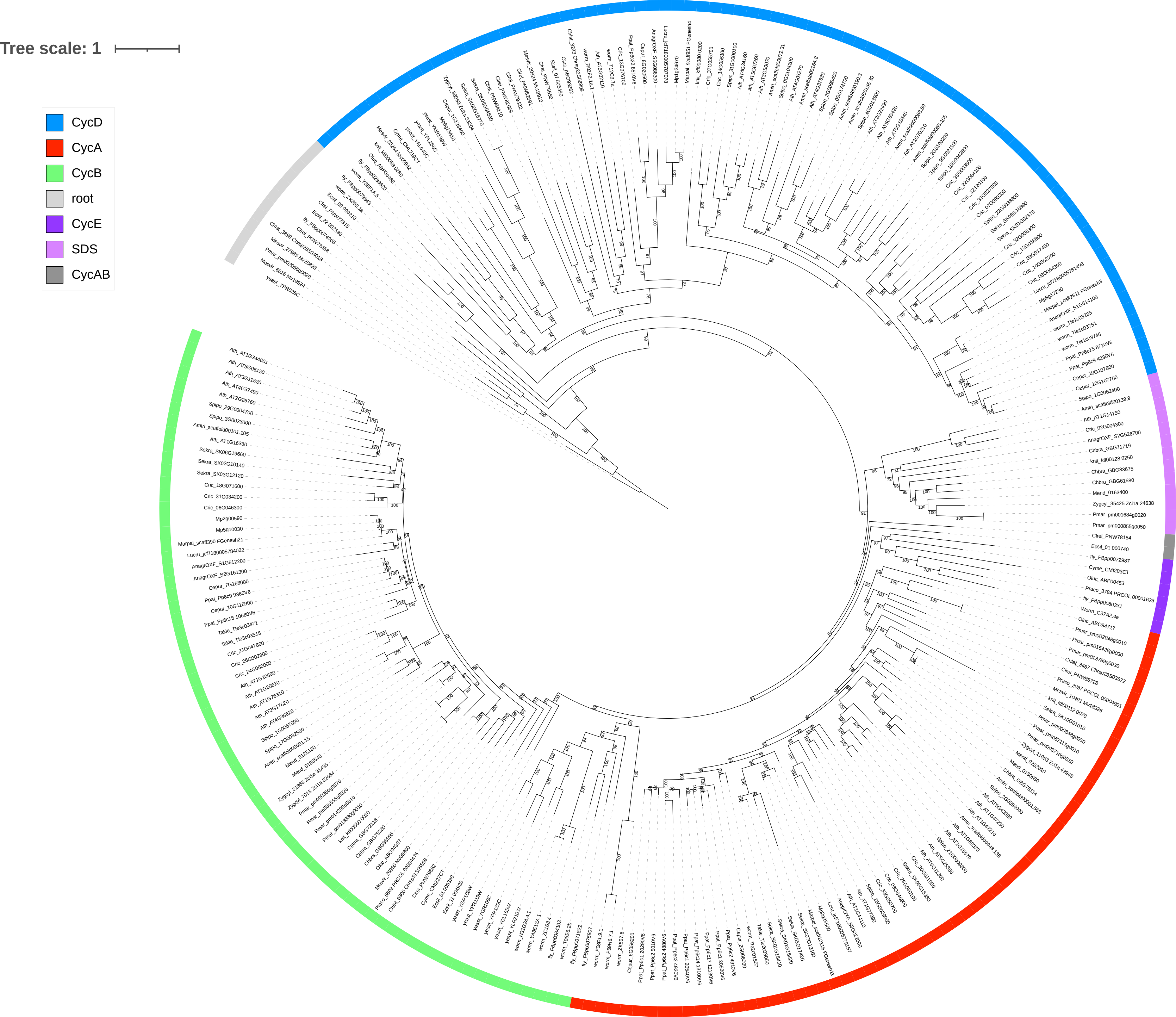
Tree scale: 1
CycD
CycA
CycB
root
CycE
SDS
CycAB

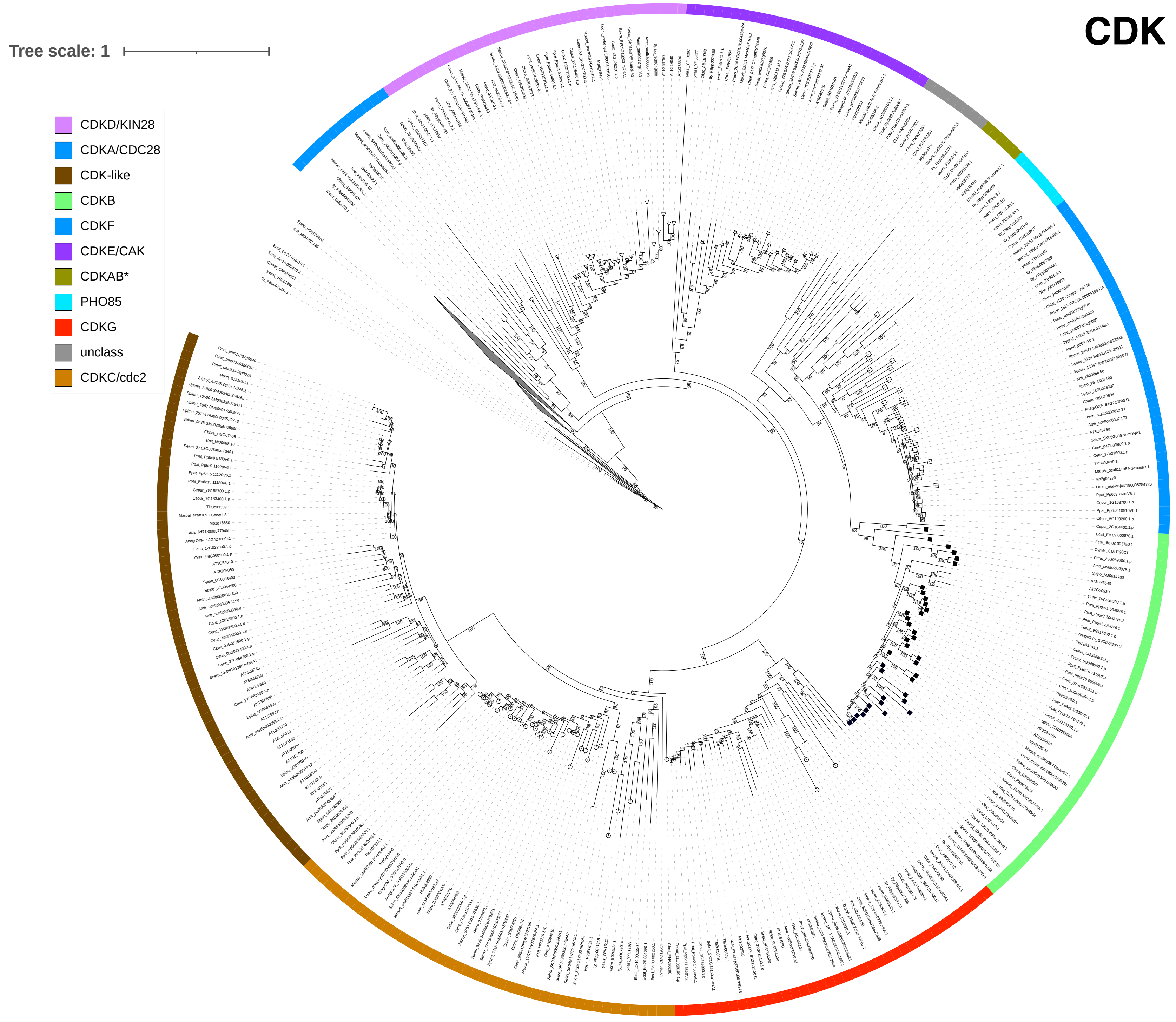
CDK
Tree scale: 1
CDKD/KIN28
CDKA/CDC28
CDK-like
CDKB
CDKF
CDKE/CAK
CDKAB*
PHO85
CDKG
unclass
CDKC/cdc2

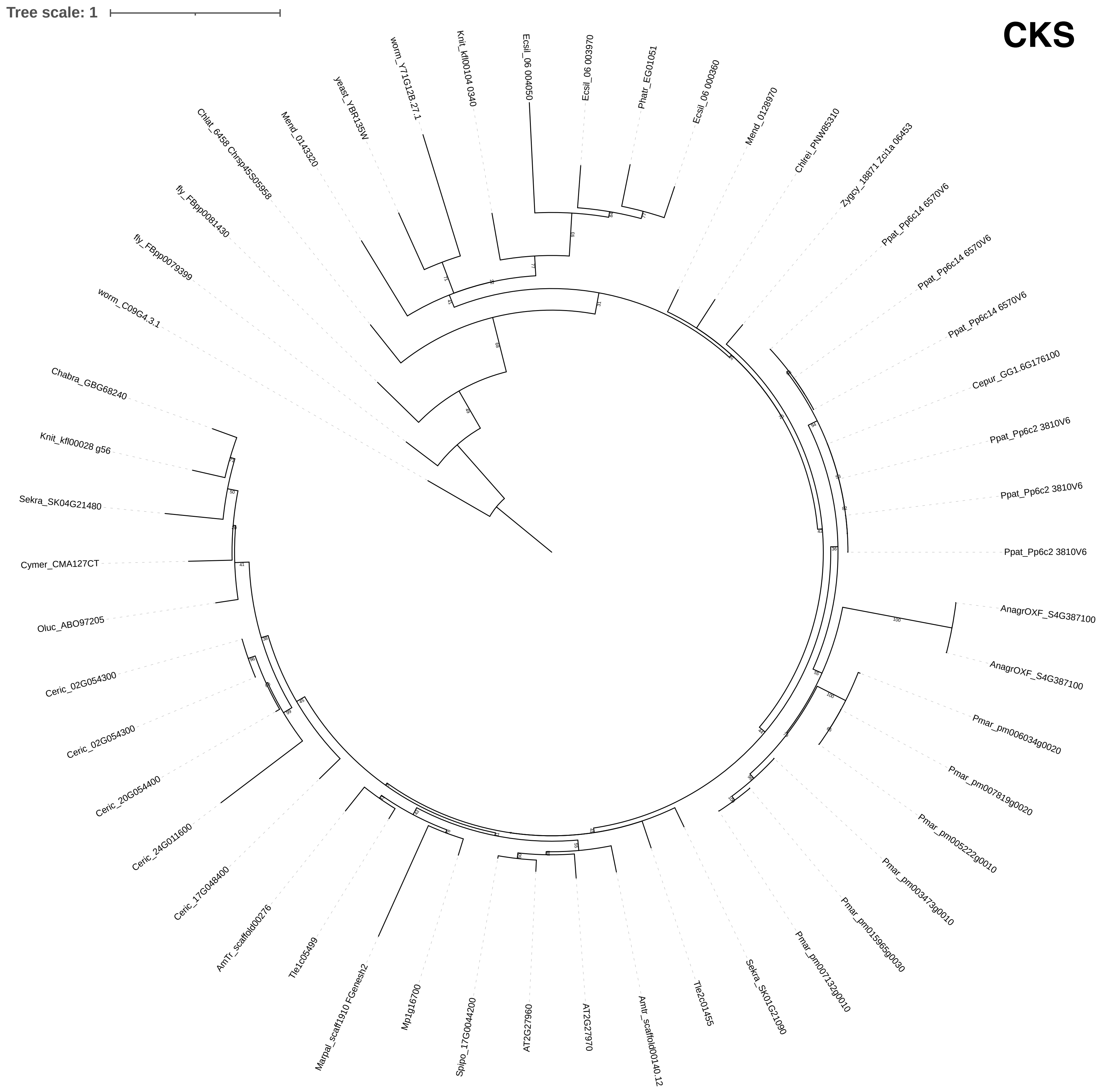
Tree scale: 1
CKS
Ecsil_06 004050
Ecsil_06 003970
Phatr_EG01051
Ecsil_06 000360
Mend_0128970
Chlrei_PNW85310
Zygcy_18871 Zci1a 06453
Ppat_Pp6c14 6570V6
Ppat_Pp6c14 6570V6
Ppat_Pp6c14 6570V6
Cepur_GG1.6G176100
Ppat_Pp6c2 3810V6
Ppat_Pp6c2 3810V6
Ppat_Pp6c2 3810V6
AnagrOXF_S4G387100
AnagrOXF_S4G387100
Pmar_pm006034g0020
Pmar_pm007819g0020
Pmar_pm005222g0010
Pmar_pm003473g0010
Pmar_pm015965g0030
Pmar_pm007132g0010
Sekra_SK01G21090
Tle2c01455
Amtr_scaffold00140.12
AT2G27970
AT2G27960
Spipo_17G0044200
Mp1g16700
Marpal_scaff1910 FGenesh2
Tle1c05499
AmTr_scaffold00276
Ceric_17G048400
Ceric_24G011600
Ceric_20G054400
Ceric_02G054300
Ceric_02G054300
Oluc_ABO97205
Cymer_CMA127CT
Sekra_SK04G21480
Knit_kfl00028 g56
Chabra_GBG68240
worm_C09G4.3.1
fly_FBpp0079399
fly_FBpp0081430
Chlat_6458 Chrsp45S05958
Mend_0143320
yeast_YBR135W
worm_Y71G12B.27.1
Knit_kfl00104 0340

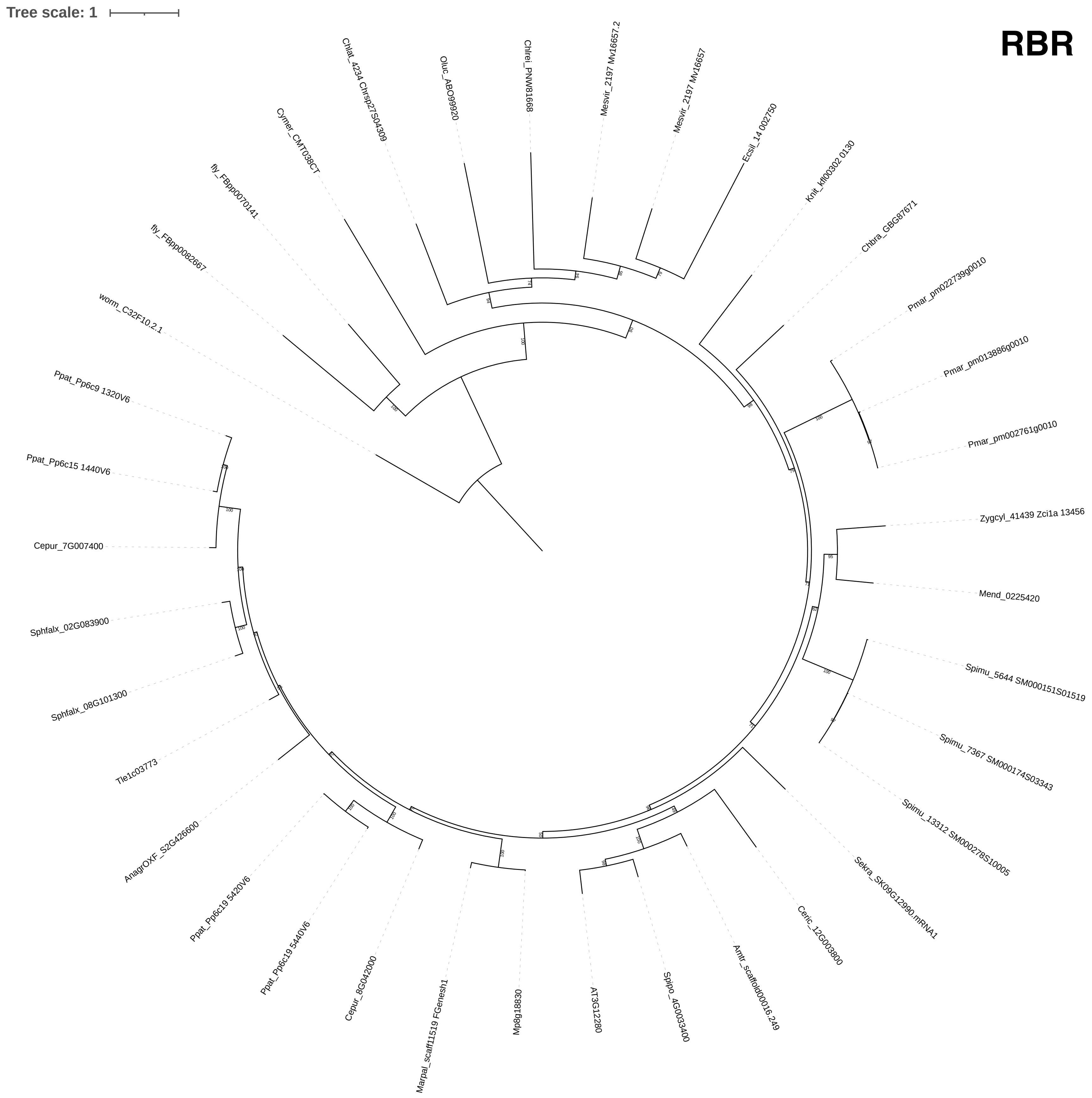
Tree scale: 1
RBR
Chlat_4234 Chrsp27S04309
Oluc_ABO99920
Chlrei_PNW81668
Mesvir_2197 Mv16657.2
Mesvir_2197 Mv16657
Ecsil_14 002750
Knit_kfl00302 0130
Chbra_GBG87671
Pmar_pm022739g0010
Pmar_pm013886g0010
Pmar_pm002761g0010
Zygcyl_41439 Zci1a 13456
Mend_0225420
Spimu_5644 SM000151S01519
Spimu_7367 SM000174S03343
Spimu_13312 SM000278S10005
Sekra_SK09G12990.mRNA1
Ceric_12G003800
Amtr_scaffold00016.249
Spipo_4G0033400
AT3G12280
Mp8g18830
Marpal_scaff11519 FGenesh1
Cepur_8G042000
Ppat_Pp6c19 5440V6
Ppat_Pp6c19 5420V6
AnagrOXF_S2G426600
Tle1c03773
Sphfalx_08G101300
Sphfalx_02G083900
Cepur_7G007400
Ppat_Pp6c15 1440V6
Ppat_Pp6c9 1320V6
worm_C32F10.2.1
fly_FBpp0082667
fly_FBpp0070141
Cymer_CMT038CT

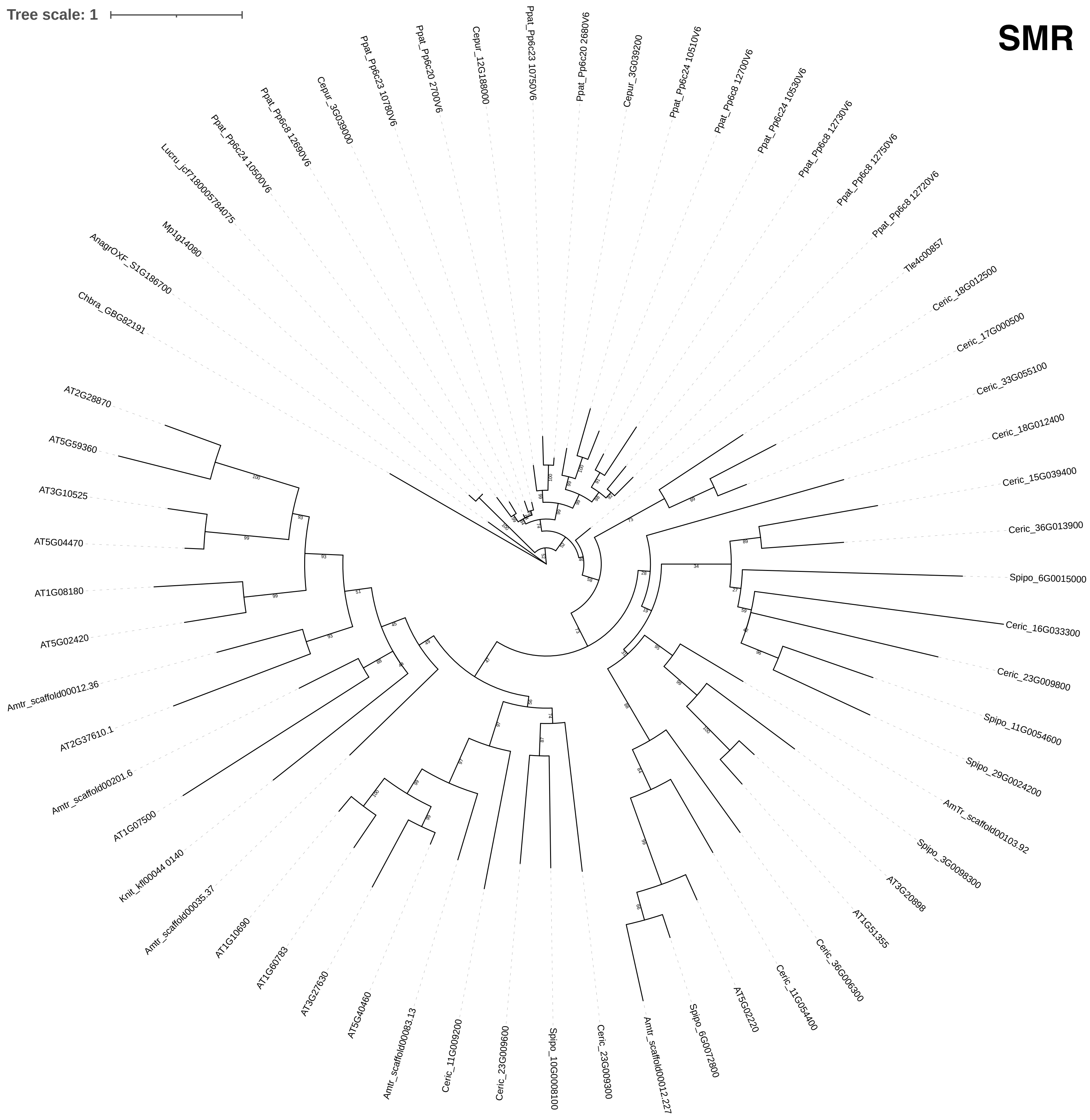

Tree scale: 1
SMR
AT2G28870
AT5G59360
AT3G10525
AT5G04470
AT1G08180
AT5G02420
Amtr_scaffold00012.36
AT2G37610.1
Amtr_scaffold00201.6
AT1G07500
Knit_kfl00044 0140
Amtr_scaffold00035.37
AT1G10690
AT1G60783
AT3G27630
AT5G40460
Amtr_scaffold00083.13
Ceric_11G009200
Ceric_23G009600
Spipo_10G0008100
Ceric_23G009300
Amtr_scaffold00012.227
Spipo_6G0072800
AT5G02220
Ceric_11G054400
Ceric_36G006300
AT1G51355
AT3G20898
Spipo_3G0098300
AmTr_scaffold00103.92
Spipo_29G0024200
Spipo_11G0054600
Ceric_23G009800
Ceric_16G033300
Spipo_6G0015000
Ceric_36G013900
Ceric_15G039400
Ceric_18G012400
Ceric_33G055100
Ceric_17G000500
Ceric_18G012500
Tle4c00857
Ppat_Pp6c8 12720V6
Ppat_Pp6c8 12750V6
Ppat_Pp6c8 12730V6
Ppat_Pp6c24 10530V6
Ppat_Pp6c8 12700V6
Ppat_Pp6c24 10510V6
Cepur_3G039200
Ppat_Pp6c20 2680V6
Ppat_Pp6c23 10750V6
Cepur_12G188000
Ppat_Pp6c20 2700V6
Ppat_Pp6c23 10780V6
Cepur_3G039000
Ppat_Pp6c8 12690V6
Ppat_Pp6c24 10500V6
Lucru_jcf7180005784075
Mp1g14080
AnagrOXF_S1G186700
Chbra_GBG82191

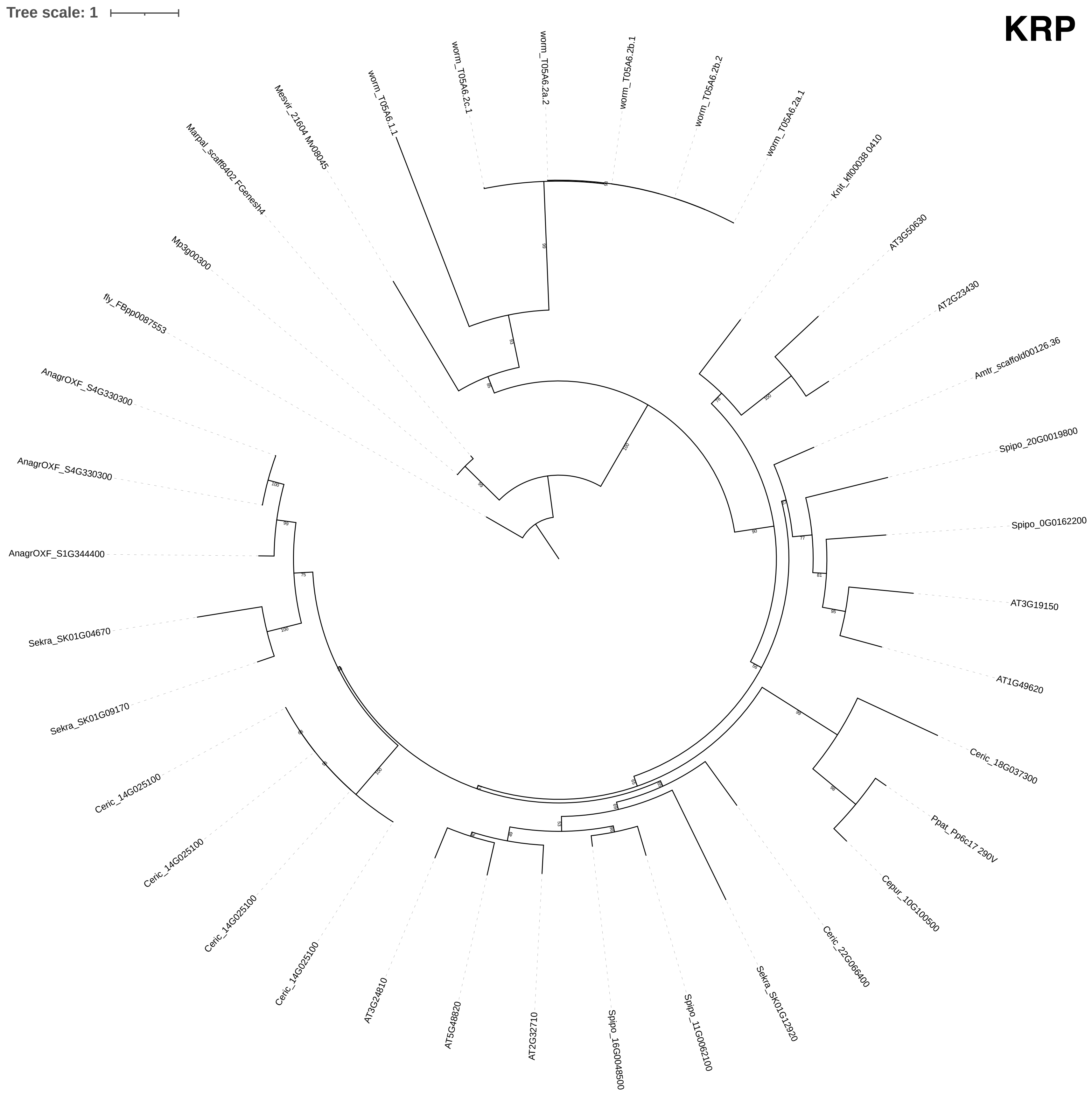

Tree scale: 1
KRP
worm_T05A6.2c.1
worm_T05A6.2a.2
worm_T05A6.2b.1
worm_T05A6.2b.2
worm_T05A6.2a.1
worm_T05A6.1.1
Mesvir_21604 Mv08045
Marpal_scaff8402 FGenesh4
Mp3g00300
fly_FBpp0087553
AnagrOXF_S4G330300
AnagrOXF_S4G330300
AnagrOXF_S1G344400
Sekra_SK01G04670
Sekra_SK01G09170
Ceric_14G025100
Ceric_14G025100
Ceric_14G025100
Ceric_14G025100
AT3G24810
AT5G48820
AT2G32710
Spipo_16G004500
Spipo_11G0062100
Sekra_SK01G12920
Ceric_22G066400
Cepur_10G100500
Ppat_Pp6c17 290V
Ceric_18G037300
AT1G49620
AT3G19150
Spipo_0G0162200
Spipo_20G0019800
Amtr_scaffold00126.36
AT2G23430
AT3G50630
Knit_kfl00038 0410

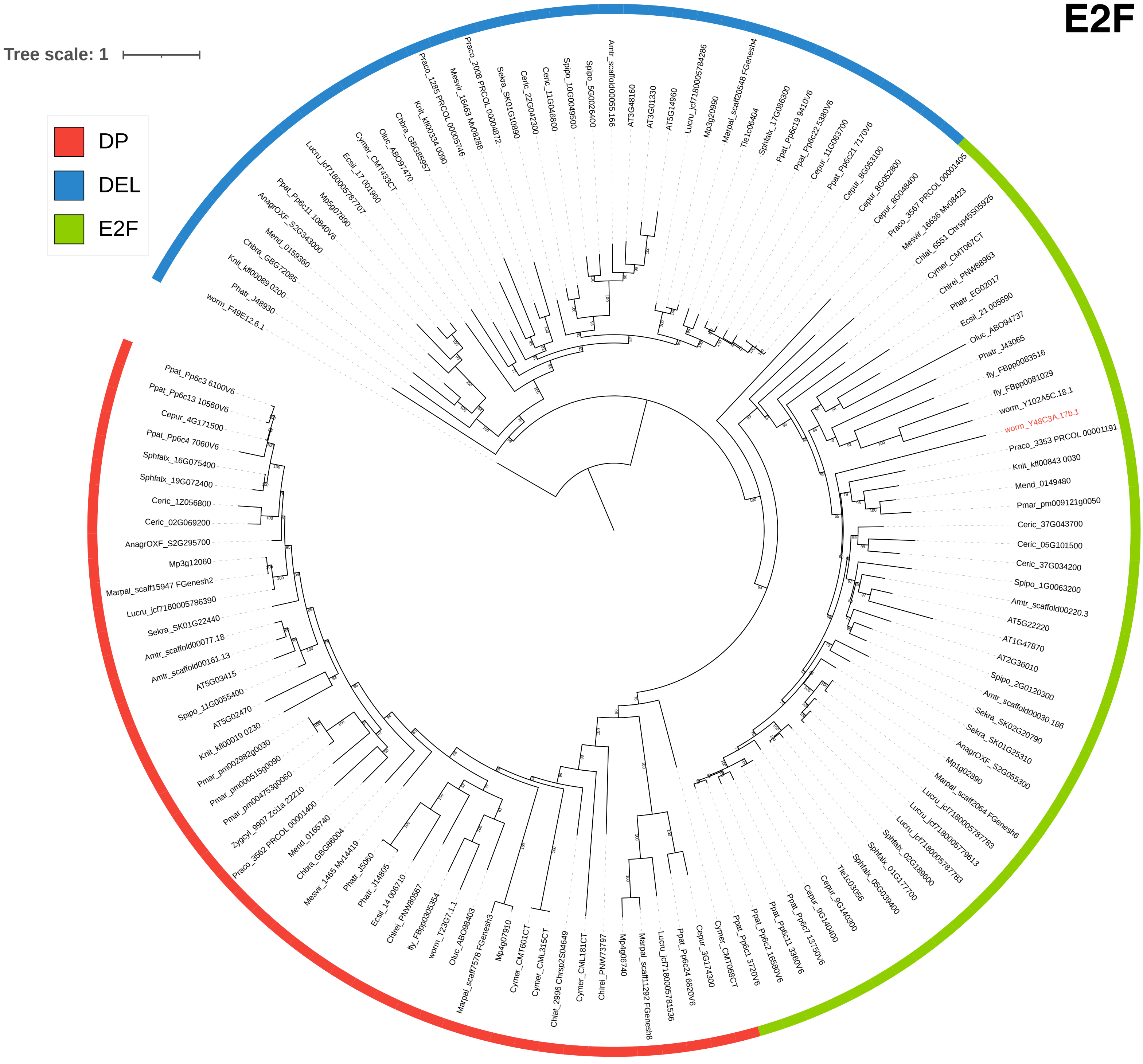
E2F
Tree scale: 1
DP
DEL
E2F
worm_F49E12.6.1
Phatr_J48930
Knit_kfl00089 0200
Chbra_GBG72085
Mend_0159360
AnagrOXF_S2G343000
Ppat_Pp6c11 10840V6
Mp5g07890
Lucru_jcf7180005787707
Ecsil_17 001960
Cymer_CMT433CT
Oluc_ABO97470
Chbra_GBG85957
Knit_kfl00334 0090
Praco_1285 PRCOL 00005746
Mesvir_16463 Mv08288
Praco_2008 PRCOL 00004872
Sekra_SK01G10890
Ceric_22G042300
Ceric_11G046800
Spipo_10G0049500
Spipo_5G0026400
Amtr_scaffold00055.166
AT3G48160
AT3G01330
AT5G14960
Lucru_jcf7180005784286
Mp3g20990
Marpal_scaff20548 FGenesh4
Tle1c06404
Sphfalx_17G086300
Ppat_Pp6c19 9410V6
Ppat_Pp6c22 5380V6
Cepur_11G083700
Ppat_Pp6c21 7170V6
Cepur_8G053100
Cepur_8G052800
Cepur_8G048400
Praco_3567 PRCOL 00001405
Mesvir_16636 Mv08423
Chlat_6551 Chrsp45S05925
Cymer_CMT067CT
Chlrei_PNW88963
Phatr_EG02017
Ecsil_21 005690
Oluc_ABO94737
Phatr_J43065
fly_FBpp0083516
fly_FBpp0081029
worm_Y102A5C.18.1
worm_Y48C3A.17b.1
Praco_3353 PRCOL 00001191
Knit_kfl00843 0030
Mend_0149480
Pmar_pm009121g0050
Ceric_37G043700
Ceric_05G101500
Ceric_37G034200
Spipo_1G0063200
Amtr_scaffold00220.3
AT5G22220
AT1G47870
AT2G36010
Spipo_2G0120300
Amtr_scaffold00030.186
Sekra_SK02G20790
Sekra_SK01G25310
AnagrOXF_S2G055300
Mp1g02890
Marpal_scaff2064 FGenesh6
Lucru_jcf7180005787783
Lucru_jcf7180005779613
Lucru_jcf7180005787783
Sphfalx_02G189600
Sphfalx_01G177700
Sphfalx_05G039400
Tle1c03056
Cepur_9G140300
Cepur_9G140400
Ppat_Pp6c7 13750V6
Ppat_Pp6c11 3360V6
Ppat_Pp6c2 16580V6
Ppat_Pp6c1 3720V6
Cymer_CMT068CT
Cepur_3G174300
Ppat_Pp6c24 6820V6
Lucru_jcf7180005781536
Marpal_scaff11292 FGenesh8
Mp4g06740
Chlrei_PNW73797
Cymer_CML181CT
Chlat_2996 Chrsp2S04649
Cymer_CML315CT
Cymer_CMT601CT
Mp4g07910
Marpal_scaff7578 FGenesh3
Oluc_ABO98403
worm_T23G7.1.1
fly_FBpp0305354
Chlrei_PNW80567
Ecsil_14 006710
Phatr_J14805
Phatr_J5060
Mesvir_1465 Mv14419
Chbra_GBG86004
Mend_0165740
Praco_3562 PRCOL 00001400
Zygcyl_9907 Zci1a 22210
Pmar_pm004753g0060
Pmar_pm000515g0090
Pmar_pm002982g0030
Knit_kfl00019 0230
AT5G02470
Spipo_11G0055400
AT5G03415
Amtr_scaffold00161.13
Amtr_scaffold00077.18
Sekra_SK01G22440
Lucru_jcf7180005786390
Marpal_scaff15947 FGenesh2
Mp3g12060
AnagrOXF_S2G295700
Ceric_02G069200
Ceric_1Z056800
Sphfalx_19G072400
Sphfalx_16G075400
Ppat_Pp6c4 7060V6
Cepur_4G171500
Ppat_Pp6c13 10560V6
Ppat_Pp6c3 6100V6

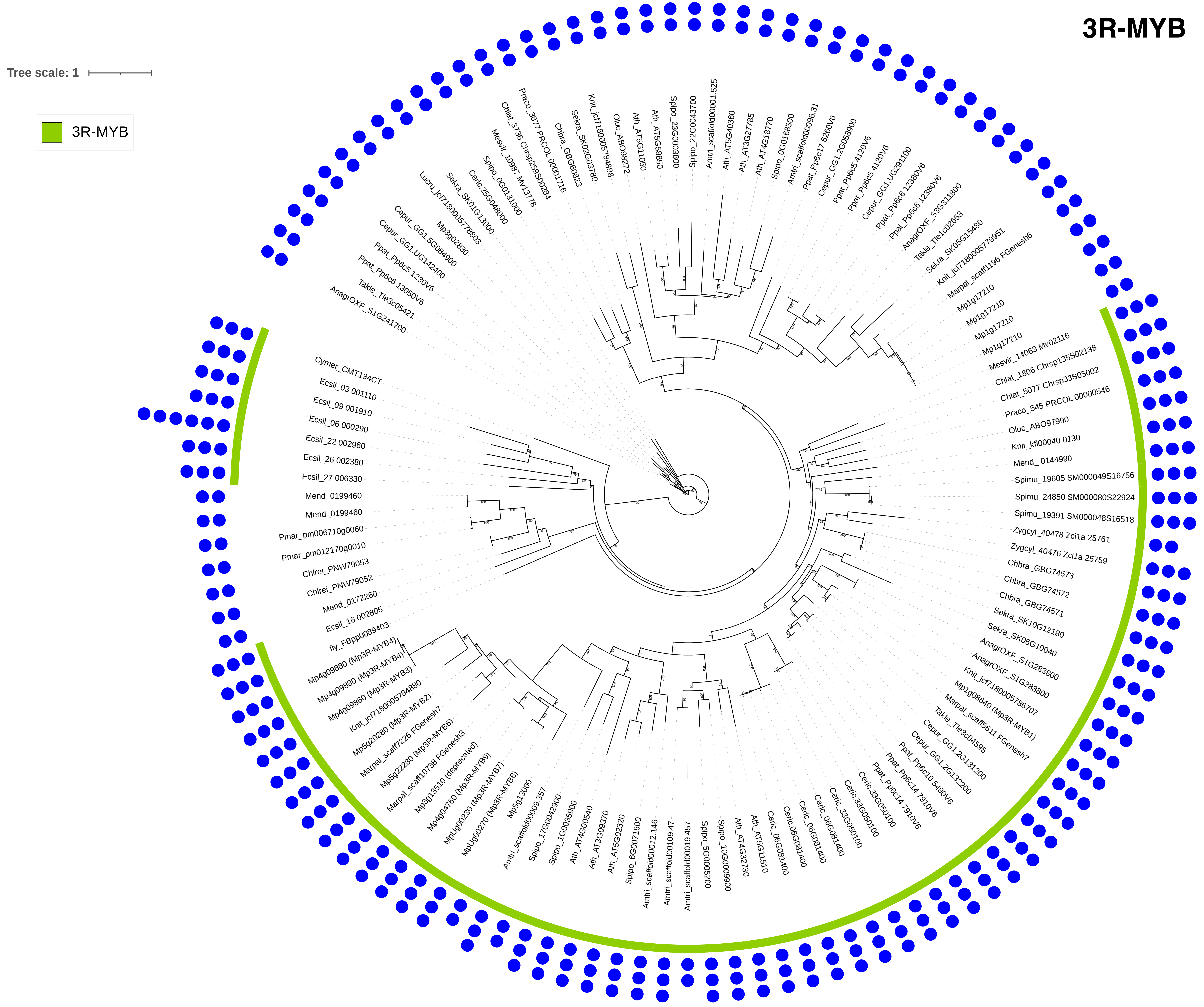
3R-MYB
Tree scale: 1
3R-MYB
Cymer_CMT134CT
Ecsil_03 001110
Ecsil_09 001910
Ecsil_06 000290
Ecsil_22 002960
Ecsil_26 002380
Ecsil_27 006330
Mend_0199460
Mend_0199460
Pmar_pm006710g0060
Pmar_pm012170g0010
Chlrei_PNW79053
Chlrei_PNW79052
Mend_0172260
Ecsil_16 002805
fly_FBpp0089403
Mp4g09880 (Mp3R-MYB4)
Mp4g09880 (Mp3R-MYB4)
Mp4g09860 (Mp3R-MYB3)
Knit_jcf7180005784880
Mp5g20280 (Mp3R-MYB2)
Marpal_scaff7226 FGenesh7
Mp5g22280 (Mp3R-MYB6)
Marpal_scaff10738 FGenesh3
Mp3g13510 (deprecated)
Mp4g04760 (Mp3R-MYB7)
MpUg00230 (Mp3R-MYB9)
MpUg00270 (Mp3R-MYB8)
Mp5g13060
Amtri_scaffold00009.357
Spipo_17G0042900
Spipo_1G0035900
Ath_AT4G00540
Ath_AT3G09370
Ath_AT5G02320
Spipo_6G0071600
Amtri_scaffold00012.146
Amtri_scaffold00109.47
Amtri_scaffold00019.457
Spipo_5G0005200
Spipo_10G0009900
Ath_AT4G32730
Ath_AT5G11510
Ceric.06G081400
Ceric.06G081400
Ceric.06G081400
Ceric.06G081400
Ceric.33G050100
Ceric.33G050100
Ppat_Pp6c14 7910V6
Ppat_Pp6c14 7910V6
Ppat_Pp6c10 5490V6
Cepur_GG1.2G132200
Cepur_GG1.2G131200
Takle_Tle3c04595
Marpal_scaff5611 FGenesh7
Mp1g08640 (Mp3R-MYB1)
Knit_jcf7180005786707
AnagrOXF_S1G283800
AnagrOXF_S1G283800
Sekra_SK06G10040
Sekra_SK10G12180
Chbra_GBG74571
Chbra_GBG74572
Chbra_GBG74573
Zygcyl_40476 Zci1a 25759
Zygcyl_40478 Zci1a 25761
Spimu_19391 SM000048S16518
Spimu_24850 SM000080S22924
Spimu_19605 SM000049S16756
Mend_0144990
Knit_kfl00040 0130
Oluc_ABO97990
Praco_545 PRCOL 00000546
Chlat_5077 Chrsp33S05002
Chlat_1806 Chrsp135S02138
Mesvir_14063 Mv02116
Mp1g17210
Mp1g17210
Mp1g17210
Mp1g17210
Marpal_scaff1196 FGenesh6
Knit_jcf7180005779951
Sekra_SK05G15480
Takle_Tle1c02653
AnagrOXF_S3G311800
Ppat_Pp6c6 12380V6
Ppat_Pp6c6 12380V6
Cepur_GG1.UG291100
Ppat_Pp6c5 4120V6
Ppat_Pp6c5 4120V6
Cepur_GG1.2G058900
Ppat_Pp6c17 6260V6
Amtri_scaffold00096.31
Spipo_0G0168500
Ath_AT4G18770
Ath_AT3G27785
Ath_AT5G40360
Amtri_scaffold00001.525
Spipo_22G0043700
Spipo_23G0003800
Ath_AT5G58850
Ath_AT5G11050
Oluc_ABO98272
Knit_jcf7180005784898
Sekra_SK02G03780
Chbra_GBG60823
Praco_3877 PRCOL 00001716
Chlat_3736 Chrsp259S00284
Mesvir_10987 Mv13778
Spipo_0G0131000
Ceric.25G048000
Sekra_SK01G13000
Lucru_jcf7180005778803
Mp3g02830
Cepur_GG1.5G084900
Cepur_GG1.UG142400
Ppat_Pp6c5 1230V6
Ppat_Pp6c6 13050V6
Takle_Tle3c05421
AnagrOXF_S1G241700
